# Trophoblast glycoprotein is required for efficient synaptic vesicle exocytosis from retinal rod bipolar cells

**DOI:** 10.1101/2022.07.25.501380

**Authors:** Colin M Wakeham, Qing Shi, Gaoying Ren, Tammie L Haley, Robert M Duvoisin, Henrique von Gersdorff, Catherine W Morgans

**Author notes:** Corresponding Author Catherine W Morgans Chemical Physiology & Biochemistry, L-334 Oregon Health & Science University 3181 SW Sam Jackson Park Rd Portland, OR 97239. **Author contributions** Study design: CMW, RD, HvG, CWM; Experimentation: CMW, HS, GR, TLH; Data analysis: CMW, HS, RD, HvG, CWM; Figure and manuscript preparation: CMW, HS, RD, HvG, CWM; Editing and review: CMW, RD, HvG, CWM.

## Abstract

Rod bipolar cells (RBCs) faithfully transmit light-driven signals from rod photoreceptors in the outer retina to third order neurons in the inner retina. Recently, significant work has focused on the role of leucine-rich repeat (LRR) proteins in synaptic development and signal transduction at RBC synapses. We previously identified trophoblast glycoprotein (TPBG) as a novel transmembrane LRR protein localized to the dendrites and axon terminals of RBCs. We now examine the effects on RBC physiology and retinal processing of TPBG genetic knockout in mice using immunofluorescence and electrophysiological techniques. The scotopic electroretinogram showed a modest increase in the b-wave and a marked attenuation in oscillatory potentials in the TPBG knockout. No effect of TPBG knockout was observed on the RBC dendritic morphology, TRPM1 currents, or RBC excitability. Because scotopic oscillatory potentials primarily reflect RBC-driven rhythmic activity of the inner retina, we investigated the contribution of TPBG to downstream transmission from RBCs to third-order neurons. Using electron microscopy, we found shorter synaptic ribbons in TPBG knockout axon terminals in RBCs. Time-resolved capacitance measurements indicated that TPBG knockout reduces synaptic vesicle exocytosis and subsequent GABAergic reciprocal feedback without altering voltage-gated Ca^2+^ currents. Thus, TPBG is required for normal synaptic ribbon development and efficient neurotransmitter release from RBCs to downstream cells. Our results highlight a novel synaptic role for TPBG at RBC ribbon synapses and support further examination into the mechanisms by which TPBG regulates RBC physiology.

**Graphical abstract:** 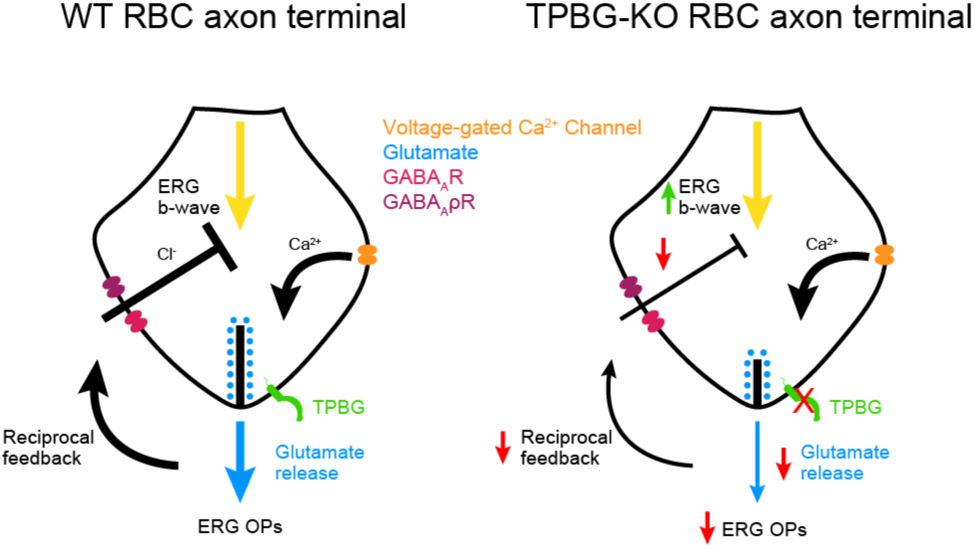

## Introduction

Rod bipolar cells (RBCs) receive light-dependent synaptic input from rod photoreceptors in the outer plexiform layer (OPL) and drive retinal output by synapsing with AII amacrine cells (AII-ACs) in the inner plexiform layer (IPL). As the only dedicated bipolar cell in the primary rod visual pathway, RBCs must be able to reliably transmit visual signals over the whole range of rod sensitivity. Accurate transmission of light signals requires precise synaptic targeting and tight control of signal transduction between RBCs and its synaptic partners. Identifying and characterizing the proteins required for the optimization and regulation of RBC synaptic development and function is essential for expanding our understanding of how RBCs contribute to vision.

Leucine-rich repeat (LRR) proteins form a large class of transmembrane proteins, each containing multiple leucine-rich motifs arranged to form extracellular protein binding domains (Kobe & Deisenhofer, 1994; Kobe & Kajava, 2001). Several LRR proteins identified in RBCs have been implicated in the development and maintenance of synaptic morphology and function. At synapses between photoreceptors and ON-bipolar cells, LRIT3 and nyctalopin are required for the localization of signal transduction components to the post-synaptic compartment (Cao, Posokhova, & Martemyanov, 2011; Hasan et al., 2019, 2020; Neuillé et al., 2017, 2015; Pearring et al., 2011). In RBC dendrites, LRRTM4 forms a trans-synaptic complex with rod spherules (Agosto & Wensel, 2020), whereas in RBC axon terminals, LRRMT4 is required for functional GABA receptor clustering, reciprocal feedback inhibition, and dyad synapse formation (Sinha et al., 2020). The ELFN family of LRR proteins form trans-synaptic complexes between photoreceptors and bipolar cells without which functioning connections fail to develop and retinal circuitry is mis-wired (Cao et al., 2015, 2020).

We recently identified trophoblast glycoprotein (TPBG, also called 5T4 or WAIF1 (Wnt-activated inhibitory factor 1)) in a phosphoproteomics screen as a novel LRR transmembrane glycoprotein in mouse RBCs that undergoes PKCα-dependent phosphorylation (Wakeham et al., 2019). TPBG contains eight N-terminal extracellular LRR motifs and a short intracellular C-terminal tail with a PDZ-interacting domain (Zhao, Malinauskas, Harlos, & Jones, 2014). TPBG localizes to the dendrites and synaptic terminals of RBCs and its expression is closely linked to eye opening and TRPM1 expression, suggesting that TPBG may play a role in RBC development or physiology (Wakeham, Ren, & Morgans, 2020).

Because the role of TPBG in the retina has not been established, we sought to characterize the effects of genetic knockout of TPBG in mouse RBCs using immunofluorescence confocal microscopy, electron microscopy, and electrophysiological techniques. RBC synapses form specialized active zones, each containing a synaptic ribbon, that permit a greater bandwidth of information transfer than conventional synapses. Ribbon synapses are noted for tight vesicle-Ca^2+^ coupling and a large supply of readily-available synaptic vesicles tethered close to the active zone (Dieck & Brandstätter, 2006; Morgans, 2000; Moser, Grabner, & Schmitz, 2020). In this study, we report that TPBG is required for normal synaptic ribbon morphology and efficient synaptic vesicle exocytosis at RBC synapses, implicating TPBG in ribbon synapse development and function.

## Methods

### Animals and statistical analysis

Constitutive TPBG knockout (TPBG-KO; KO) mice were B6;129S5-Tpbg^tm1Lex^/Mmucd (MMRRC; UC Davis; Davis, CA, USA; Cat# 031630-UCD; RRID: MMRRC_031630-UCD). KO mice were purchased as cryopreserved sperm and reconstituted into a C57BL/6J (The Jackson Laboratory; Bar Harbor, M; Cat# 000664; RRID: IMSR_JAX:000664) background. Age-matched homozygous wild type (WT) littermates were used as controls. We used mice of both sexes and all mice were maintained on a 12-hour light/dark cycle and provided food and water ad libitum. Animals for immunofluorescence and electroretinogram experiments were 3-6 months old while animals used for patch clamp electrophysiology experiments were three months old. At least three mice were used from each condition for each experiment except for electron microscopy, which used one block per genotype. All animal procedures were in accordance with the National Institutes of Health guidelines and approved by the Oregon Health & Science University Institutional Animal Care and Use Committee.

Statistical analyses were performed in Prism 9 (GraphPad; San Diego, CA, USA) and reported as mean ± SEM. We detected significance between conditions using unpaired Welch’s *t*-tests and Brown-Forsythe ANOVA tests. Statistical significance is noted by asterisks (not significant (ns): p > 0.05; *: p < 0.05; **: p < 0.01; ***: p < 0.001; ****: p < 0.0001).

### Immunofluorescence

Fresh eyecups were prepared from dissected eyes by cutting behind the ora serrata and removing the cornea and lens. For sectioning, eyecups were fixed for 30 min by immersion in fresh 4% paraformaldehyde (PFA) in PBS. The fixed eyecups were washed in PBS and then cryoprotected via sequential immersion in 10, 20, and 30% sucrose in PBS. Sections were cut at 20 μm thickness. Retina sections were post-fixed for 10 minutes in 4% PFA, blocked and permeabilized by incubation at room temperature for one hour in antibody incubation solution (AIS; 3% normal horse serum, 0.5% Triton X-100, 0.025% NaN_3_ in PBS), and then incubated in primary antibodies diluted in AIS for one hour at room temperature. After washing with PBS, the sections were incubated in secondary antibodies diluted in AIS for one hour at room temperature. The slides were washed again in PBS and then mounted with VECTASHIELD PLUS antifade mounting media with DAPI (Vector Laboratories; Cat# H-2000-10).

For whole mount immunofluorescence, eyecups were fixed for 45 min by immersion in fresh 4% PFA in PBS. The fixed eyecups were washed in PBS then retinas were removed prior to blocking and permeabilization for one hour in AIS. The retinas were incubated in primary antibodies diluted in AIS for four days at 4° C then washed overnight in PBST (PBS + 0.05% Triton X-100) at 4° C. Retinas were incubated in secondary antibodies diluted in AIS overnight at 4° C then washed again in PBST. Finally, the retinas were mounted with VECTASHIELD PLUS antifade mounting media.

Immunofluorescence images were taken with a Leica TCS SP8 X confocal microscope (Leica Biosystems; Wetzlar, Germany) using a Leica HC PL APO CS2 63x/1.40 oil immersion objective (Leica; Cat# 15506350). Brightness and contrast were adjusted equally across genotypes using Fiji (Schindelin et al., 2012). A Gaussian blur using a radius of 0.5 pixels was used to remove graininess from images. See figure legends for specific information on Z-step distance and number of steps. To compare the spatial relationship between presynaptic and postsynaptic proteins in the OPL, ROI lines were drawn across Ca_v_1.4-GPR179 pairs and the intensity of both fluorescent channels was plotted against the line length in Fiji. The distance between the two fluorescence peaks approximates the distance across the synapse. To examine fluorescent puncta in RBC axon terminals in the IPL, PKCα binaries were created in Fiji using default auto-thresholds to isolate RBC axon terminals. These binaries were then merged with the other channels to remove PKCα-negative puncta. Fluorescent puncta were counted in PKCα-positive axon terminals and normalized to axon terminal area. Mander’s colocalization coefficients were calculated using the JACoP plugin in Fiji to compare colocalization patterns between Ca_v_1.4 and RIBEYE puncta after thresholding with Otsu’s method. The M1 coefficient was used for the fraction of Ca_v_1.4 containing RIBEYE and the M2 coefficient was for the fraction of RIBEYE containing Ca_v_1.4. Colocalization of puncta in PKCα-negative areas was also calculated as a control.

### Immunoblotting

Retinas were extracted from freshly dissected eyes, suspended in chilled lysis buffer (50 mM Tris pH 7.4, 150 mM NaCl, 1 mM EDTA, 1% Triton X-100, 1% deoxycholate, 0.1% SDS) with 1X protease/phosphatase inhibitor cocktail (Cell Signaling Technology; Danvers, MA, USA; Cat# 5872) and homogenized with a Teflon-glass homogenizer. The lysate was centrifuged for 15 minutes at 16,400 rpm and 4° C, and the pellet was discarded. Retinal lysates were diluted to a protein concentration of 1 μg/μl in lysis buffer and brought to 1X NuPAGE LDS Sample Buffer (Thermo Fisher Scientific; Cat# NP0007) and 1X NuPAGE Sample Reducing Agent (Thermo Fisher Scientific; Cat# NP0009). Pre-cast NuPAGE 1 mm 4-12% Bis-Tris gels (Thermo Fisher Scientific; Cat# NP0322BOX) were loaded and run at 200 V and 140 mA for 55 minutes in 1X NuPAGE BOLT SDS Running Buffer (Thermo Fisher Scientific; Cat# B0001).

Proteins were transferred onto PVDF membranes using a wet transfer system and 1X NuPAGE Transfer Buffer (Thermo Fisher Scientific; Cat# NP00061) with 5% MeOH at 300 mA for two hours. The membranes were then rinsed with methanol and blocked for one hour in Odyssey Blocking Buffer TBS (LI-COR Biosciences; Cat# 927-50003) on a shaker at room temperature, before being incubated in primary antibody diluted in Odyssey buffer at 4° C overnight. The membranes were washed in TBST (Tris-buffered saline with 0.1% Tween-20), then incubated in secondary antibody diluted in Odyssey buffer for one hour at room temperature before being washed in TBST. Dried blots were imaged using a LI-COR Odyssey CLx Imaging System at 680 and 800 nm.

### Electron microscopy

Mice were euthanized and enucleated and the retinas were quicky dissected and fixed for 30 min by immersion in 3% PFA and 1% glutaraldehyde. The tissue was then processed for electron microscopy using the Dresden protocol (Paridaen, Wilsch-Bräuninger, & Huttner, 2013). Three-dimensional data was acquired using the Helios G3 NanoLab DualBeam FIB-SEM as previous described. Briefly, the resin blocks were trimmed to contain the ganglion cell layer and the IPL near the top edge of the block, mounted to a 45° pre-tilted stub, and coated with 8 nm of carbon. A focused beam of gallium atoms ablated 4 nm off the surface of the sample (FIB conditions: 30 keV accelerating voltage, 790 pA beam current) over the SEM imaging area of 25 μm x 10 μm. The freshly ablated surface was then imaged by backscattered electrons using the In-Column Detector (SEM conditions: 3 keV accelerating voltage, 400 pA beam current, 4 nm per pixel, 3 μs dwell time; image resolution: 6144 x 4086, 4 nm isotropic). Ablation and imaging cycles were run over a three-day period resulting in a sample depth of roughly 5 μm (1373 images for the WT dataset and 1284 images for the TPBG-KO dataset). Image stacks were registered using Linear Stack Alignment with the SIFT algorithm in Fiji and cropped to remove alignment artifacts. RBC synaptic terminals were identified by their morphology (size, shape, presence of synaptic ribbons, and arrangement of postsynaptic elements) and their proximity to the ganglion cell layer.

### Electroretinogram recording

Mice were dark-adapted overnight and prepared for recording under dim red light. Anesthesia was initiated via IP injection of ketamine:xylazine (100:10 mg/kg) and maintained with booster injections (30:3 mg/kg) at approximately 30 minute intervals until completion of the experiment. Body temperature was maintained at 36.5-38° C. Before recording, the pupils were dilated with 2.5% phenylephrine and 1% tropicamide, and the cornea was anesthetized with 1% topical proparacaine. A wire loop placed under the upper teeth was used to draw the mouse into a custom-made holder that stabilized the head and permitted delivery of 95% O_2_/5% CO_2_ (carbogen; ∼0.25 l/min). The recording was made using a custom contact lens electrode featuring a central platinum wire placed against the cornea with a small drop of hypromellose to prevent drying. A loop electrode placed behind the eye served as a reference electrode and a needle electrode placed in the tail served as a ground. The mice were placed in a Ganzfield sphere and light stimuli were provided by custom 502 nm LED photostimulators. Light stimulus intensity was controlled with neutral density filters and by altering flash duration and was measured post-hoc with an ILT5000 radiometer (International Light; Newburyport, MA; USA) using a scotopic filter. Traces were recorded with customized software (ERGLab; Richard Weleber; Casey Eye Institute; Portland, OR, USA).

Full-field scotopic electroretinograms (ERGs) from both eyes were recorded simultaneously after a series light flashes ranging from 8.71 x 10^-1^ to 8.51 x 10^1^ cd•s/m^2^ (-0.06 to 1.93 log(cd•s/m^2^) (Xiong et al., 2015). For dimmer intensities (-1 to 0 log(cd•s/m^2^)), three trials were averaged with inter-flash intervals of 20 seconds. For brighter intensities (0 to 2 log(cd•s/m^2^)), two trials were averaged with inter-flash intervals of 30 to 120 seconds.

ERGLab data was exported for offline processing using a custom Python script and the SciPy package (https://docs.scipy.org/doc). Statistical analyses were performed in Prism 9 (GraphPad; San Diego, CA, USA). The start of the flash stimulus was set to time zero and traces were baselined. A-waves and b-waves were isolated using a low-pass filter (60 Hz) and OPs were isolated with a Butterworth band-pass filter (30-300 Hz) and the “signal.filtfilt” function in SciPy (**Figure 2: Supplementary Figure 1**). The a-wave amplitudes were measured from baseline-to-trough, b-wave amplitudes were measured from a-wave trough-to-b-wave peak, and the b/a-wave ratio was calculated to normalize responses. OP amplitude was measured from trough-to-peak of the largest wave and the OP/b-wave ratio was calculated using b-wave amplitudes from the same recording.

### Patch clamp electrophysiology

Retinas were isolated and embedded in 3% low melting point agarose (Sigma; Cat# A0701) in carbogenated bicarbonate-buffered Ames medium (Ames & Nesbett, 1981; Ames Medium w/L-Glutamate; US Biological; Swampscott, MA, USA; Cat# A1372-25) and 200-250 μm slices were prepared with a Leica VT1200 vibrating microtome. For chemically simulated light response experiments, retinas were sliced in carbogenated Ames. For all other recordings, slicing was performed in chilled and carbogenated low-Na^+^, high-sucrose cutting solution containing (in mM): 210 sucrose, 35 NaHCO_3_, 20 HEPES, 10 glucose, 6 MgCl_2_, 3 sodium pyruvate, 2.5 KCl and 0.5 CaCl_2_ and buffered to pH 7.4 with NaOH. Retina slices were stored at room temperature in carbogenated Ames medium before being transferred to the recording chamber. Heated (31-33° C) Ames medium was continuously perfused over the retinal slices during all patch-clamp recordings.

Thick-walled borosilicate glass pipettes (World Precision Instruments; Sarasota, Fl, USA; Cat# 1B150F-4) were pulled using a Narishige PP-830 electrode puller (Tokyo, Japan). Ultrapure salts were purchased from Sigma-Aldrich (St. Louis, MO, USA). For RBC visualization, the fluorescent tracer Alexa Fluor 594 Hydrazide (100 μM; Thermo Fisher Scientific; Cat# A10438) was added to the pipettes. Whole-cell voltage-clamp and current-clamp recordings were performed in photopic conditions using a double EPC-10 patch clamp amplifier controlled by Patchmaster software (HEKA Elektronik, Lambrecht/Pfalz, Germany) or an Axon Axopatch 200B controlled by pCLAMP software (Molecular Devices; San Jose, CA, USA). Data was acquired at a 20 kHz sampling rate and filtered with a 2.9 kHz low-pass Bessel filter. RBCs were targeted visually using Dodt gradient contrast microscopy and confirmed post-hoc with cell-filling and epifluorescence microscopy. Both pCLAMP and Patchmaster data sets were exported to Igor Pro (WaveMetrics, Lake Oswego, OR, USA) for offline analysis with custom Igor scripts and statistical analysis was performed in Prism. We used a P/10 leak subtraction protocol (-80 mV to -73 mV) in voltage-clamp to isolate voltage-gated Ca^2+^ currents and GABAergic feedback currents; leak traces for each trial were baselined, summated, and subtracted from the raw pulse trace.

To record chemically simulated light responses, pipettes were pulled to 10-12 MΩ and backfilled with a potassium gluconate solution containing (in mM): 125 K-gluconate, 8 KCl, 5 HEPES, 1 MgCl_2_, 1 CaCl_2_, 3 adenosine 5’-triphosphate magnesium salt (Mg-ATP), 0.5 guanosine 5’-triphosphate disodium salt (Na_2_-GTP), and 5 EGTA, and pH was adjusted to 7.3 with KOH. The Ames medium was supplemented with 4 μM L-(+)-2-amino-4-phosphonobutyric acid (L-AP4), an mGluR6 agonist, to maintain a simulated dark-adapted state. A puffer pipette (12-18 MΩ) was maneuvered into the dendritic field of the target cell, and the cell was patched in voltage-clamp and held at -60 mV. The mGluR6 antagonist [RS]-α-cyclopropyl-4-phophophenylglycine (CPPG) was applied directly to the dendrites using a Picospritzer II (Parker Hannifin; Hollis, NH, USA; 600 μM at 20 psi) for five seconds and the RBC response was recorded. For each cell, four sweeps were recorded at 15-second intervals.

For current-clamp experiments, pipettes were pulled to 9-12 MΩ and backfilled with a potassium gluconate solution containing (in mM): 104 K-gluconate, 10 KCl, 10 HEPES, 4 Mg-ATP, 0.4 Na_2_-GTP, 10 Na_2_-Phosphocreatine, and 2 EGTA, and pH was adjusted to 7.3 with KOH. The Ames medium was supplemented with 1 mM CaCl_2_ and the synaptic blockers (in μM): 0.5 strychnine (glycine receptor blocker), 3 gabazine (SR 95531; GABA_A_ receptor blocker), 50 [1,2,5,6-tetrahydropyridin-4-yl]methylphosphinic acid (TPMPA; GABA_A_ρ receptor blocker), and 4 L-AP4 to block excitatory and inhibitory inputs. The total external CaCl_2_ concentration was estimated to be 2.15 mM. Cells were first patched in voltage-clamp and then the amplifier was switched to fast current-clamp mode using the EPC-10 Gentle CC-Switch feature. Resting membrane voltage was recorded at 0 pA and then current was injected for 100 ms from -80 pA to 80 pA in 20 pA steps. Peak voltage was measured as the peak (min or max) voltage immediately after the onset of the current pulse, and sustained voltage was measured as the average over the final 10 ms of the pulse. For each cell, five sets of sweeps were recorded at five-second intervals.

For the remaining experiments, the pipettes were pulled to 5-8 MΩ and backfilled with a cesium gluconate solution containing (in mM): 94 Cs-gluconate, 10 HEPES, 20 tetraethyl ammonium chloride (TEA-Cl), 4 Mg-ATP, 0.4 Na_2_-GTP, 10 Na_2_-Phosphocreatine, and 2 EGTA, and pH was adjusted to 7.3 with CsOH.

For voltage-clamp time-resolved capacitance recordings, the Ames medium was supplemented with 1 mM CaCl_2_ and (in μM): 0.5 strychnine, 3 gabazine, 50 TPMPA, and 4 L-AP4 to isolate voltage-gated Ca^2+^ currents and maximize membrane resistance. Pipettes were wrapped with parafilm to reduce pipette capacitance and electrical noise. Membrane capacitance (C_m_), membrane resistance (R_m_), and series resistance (R_s_) were measured using the “sine + DC” method implemented in Patchmaster software (Balakrishnan, Puthussery, Kim, Taylor, & von Gersdorff, 2015; Gillis, 2000; Oltedal & Hartveit, 2010). Exocytosis was evoked in voltage-clamp mode with a 100 ms depolarizing pulse from -80 mV to -10 mV to activate voltage-gated Ca^2+^ currents. Resting C_m_ was measured using a sinusoidal voltage command (200 Hz, 30 mV trough-to-peak) superimposed over the membrane holding potential (-80 mV) and one measurement was generated per cycle (**Figure 5A**). An increase in exocytosis (ΔC_m_) was measured from the change in C_m_ values before and after the depolarizing pulse. A baseline was established by averaging the resting C_m_ over the one second immediately preceding the voltage step. C_m_ measurements immediately after repolarization were excluded from analysis to allow for the full decay of evoked conductances and C_m_ was averaged over the following 500 ms. Cells were discarded if changes in C_m_ correlated significantly with changes in R_m_ or R_s_. For each cell, five sets of sweeps were recorded with 15 second intervals. To remove noise, C_m_ data was filtered through a 5 Hz low-pass FIR filter. When different frequency sinusoidal voltage commands were used, recordings were resampled to 200 Hz for ease of comparisons.

To record reciprocal inhibitory currents, the Ames medium was supplemented with 1 mM CaCl_2_, 4 μM L-AP4, and 0.5 μM strychnine. Feedback current was evoked in voltage-clamp mode with a 100 ms depolarizing pulse from -80 mV to -10 mV to activate voltage-gated Ca^2+^ currents and subsequent outward GABAergic inhibitory currents. We isolated inhibitory currents (I_Inhib_) by subtracting recordings with GABA blockers (I_Ca_) were from recordings without GABA blockers (I_Total_), and calculated the charge transfer across the duration of the depolarizing pulse. Five sets of sweeps were recorded at five-second intervals.

## Results

### TPBG is not required for normal gross RBC morphology

We used the constitutive Tpbg knockout mouse line B6;129S5-Tpbg^tm1Lex^/Mmucd (referred to as TPBG-KO or KO) to analyze the consequences of genetic deletion of TPBG in the retina. Wild type (WT) littermates were used as controls. Expression and localization of TPBG were examined by western blotting and immunofluorescence confocal microscopy (**Figure 1**). As previously reported (Wakeham et al., 2020), the TPBG antibody labels RBC cell bodies, dendrites, and axon terminals in the WT retina, as well as a narrow band in the middle of the IPL corresponding to the processes of TPBG-expressing amacrine cells. TPBG immunoreactivity was absent throughout the KO retina (**Figure 1A**); this was confirmed by immunoblotting (**Figure 1B**) of WT and KO retinal lysates with the TPBG antibody using β-actin as a loading control. Using an antibody against PKCα in WT and TPBG-KO retina, we found no discernable disruption in morphology of RBC dendrites or axon terminals in KO retinas (**Figure 1C-D**).

**Figure 1:**
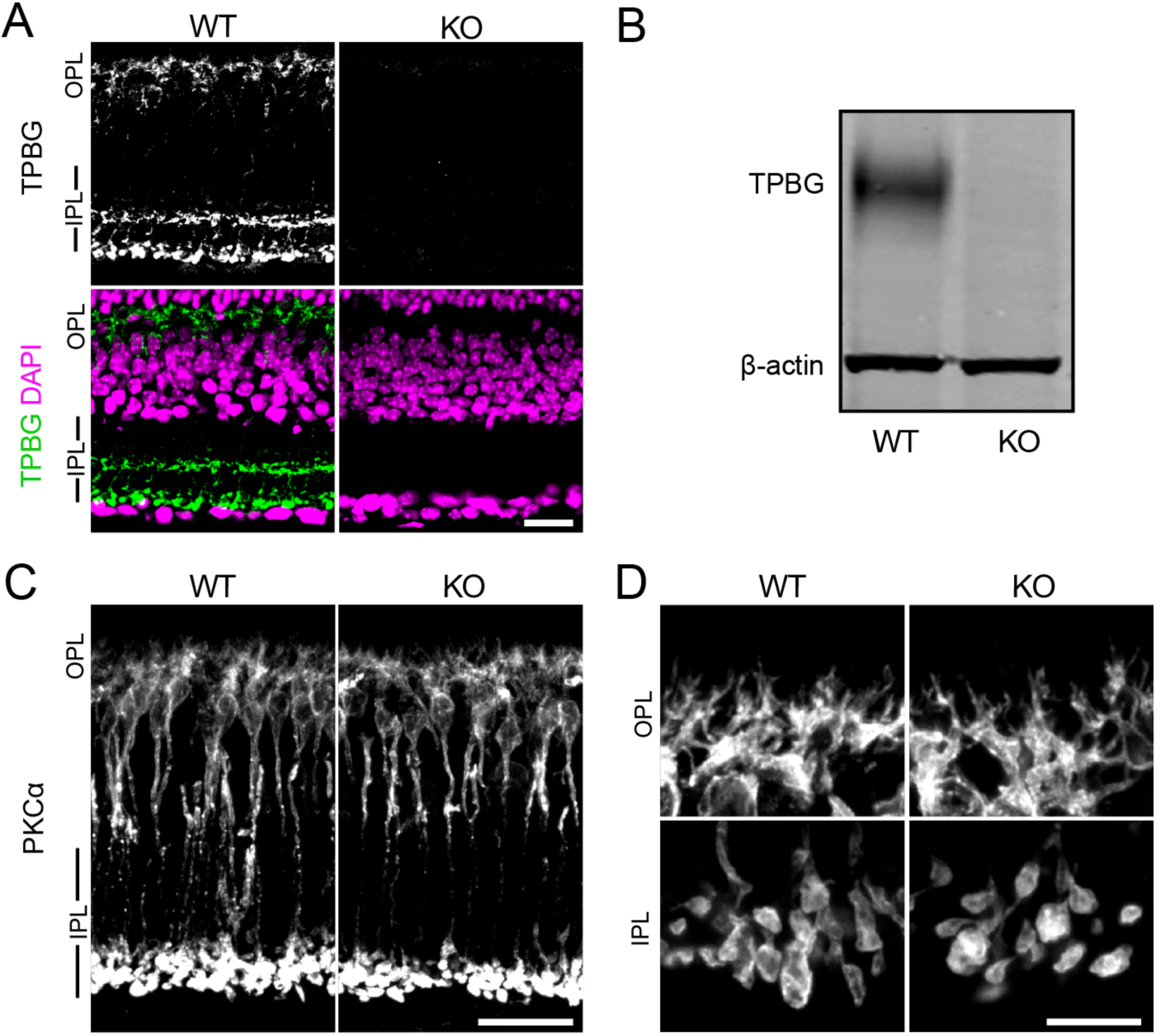
TPBG-KO does not alter gross RBC morphology. **(A)** Immunofluorescence confocal microscopy of WT (left) and TPBG-KO (KO; right) retina sections labeled with an antibody against TPBG (green). The nuclear stain DAPI (bottom; magenta) was used to highlight the neuronal layers of the retina. **(B)** Immunoblot of WT and TPBG-KO (KO) retinal lysates probed for TPBG and β-actin. The two channels were imaged separately and merged. **(C)** Confocal microscopy of WT (left) and TPBG-KO (KO; right) retina sections labeled with an antibody against PKCα. **(D)** Close-up images depicting the OPL (top) and IPL (bottom) for better visualization. For all images, Z-projections of 11 sections were used with a Z-step distance of 0.415 μm for a total thickness of 4.57 μm. Intensity and contrast was adjusted equally across genotypes. OPL: outer plexiform layer; IPL: inner plexiform layer. Scale bars: **(A and C)** 20 μm; **(D)** 10 μm.

### Genetic knockout of TPBG alters the scotopic ERG light response

The light response of RBCs is reflected in the b-wave of the dark-adapted (scotopic) full-field electroretinogram (ERG), a commonly-used technique for recording the electrical responses of the whole retina *in vivo*. Different components of the ERG waveform correspond to the activity of specific cell types in the retina. The a-wave is an initial negative deflection corresponding to hyperpolarization of the photoreceptors (Robson & Frishman, 2014), whereas the b-wave, a larger and slower positive deflection, primarily corresponds to the subsequent depolarization of the ON-bipolar cells (Stockton & Slaughter, 1989). Rod driven activity can be isolated by dark-adapting the mouse before testing; therefore, the dark-adapted ERG b-wave represents the activity of RBCs in response to light stimuli.

We recorded the scotopic ERG light responses in dark-adapted anesthetized WT (blue; *n* = 12) and TPBG-KO (KO; orange; *n* = 14) mice across several stimulus intensities, from dim (-0.06 log(cd•s/m^2^)) to bright (1.93 log(cd•s/m^2^)). We applied a 60 Hz low-pass filter to remove oscillatory potential (OP) contamination from the ERG a-and b-waves, and a 30-300 Hz Butterworth band-pass filter to isolate OPs for further analysis (**Figure 2A**). There was no difference in a-wave amplitudes across intensities (**Figure 2: Supplementary Figure**), indicating that the KO did not affect the rod photoreceptor light responses, but there was an enlarged b-wave in the TPBG-KO mouse (**Figure 2B**). Because the ERG b-wave is dependent on upstream photoreceptor activation, which is represented by the a-wave, it is common practice for ERG b-wave amplitudes to be normalized to a-wave amplitudes for comparison across animals. The TPBG-KO increased the normalized b-waves (b/a-wave ratio) across the range of stimuli intensities (**Figure 2C**), indicating a larger and more prolonged RBC light response in TPBG-KO mice.

**Figure 2:**
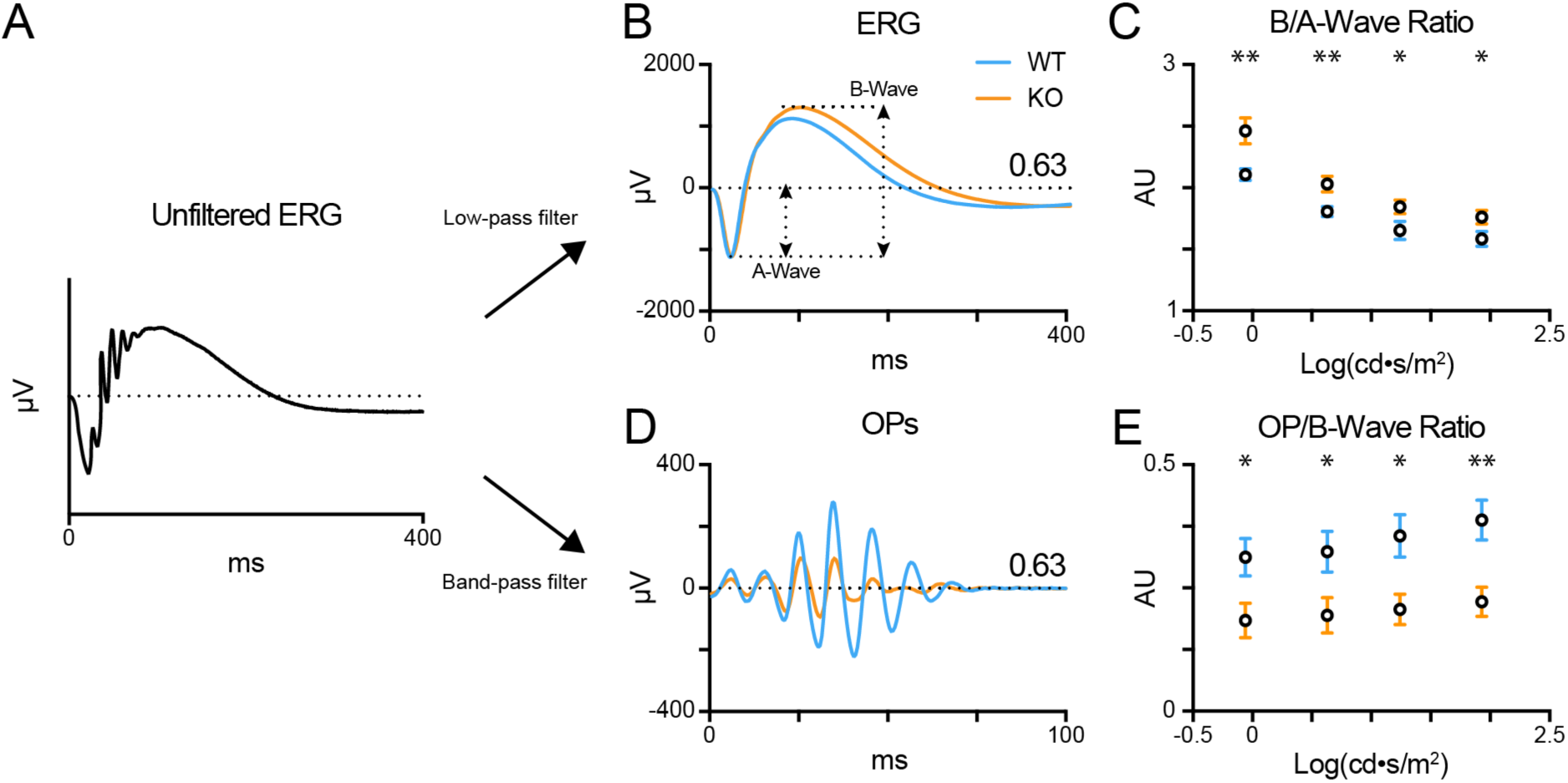
TPBG-KO alters the scotopic ERG after bright stimuli. Scotopic electroretinograms (ERGs) were recorded from dark-adapted WT (blue; *n* = 12) and TPBG-KO (KO; orange; *n* = 14) mice after light stimuli (-0.06 to 1.93 log(cd•s/m^2^)). **(A)** After baselining the raw, unfiltered ERG traces, a 60 Hz low-pass filter was used to remove OPs to better visualize the a- and b-waves. A 30-300 Hz Butteworth band-pass filter was used to isolate OPs from the ERG for quantification. **(B)** Average ERG recordings after light stimuli of 0.63 log(cd•s/m^2^). A-wave amplitudes were measured from baseline to the a-wave trough and b-wave amplitudes were measured from the a-wave trough to the b-wave peak. **(C)** Quantification of b-wave amplitudes normalized to a-wave amplitudes. **(D)** Average oscillatory potentials (OPs) after light stimuli of 0.63 log(cd•s/m^2^). **(E)** Quantification of OP amplitudes normalized to b-wave amplitudes. OP amplitudes were measured from the trough to peak of the largest wave. Quantified data is presented as the mean ± SEM. Statistical significance was determined using Welch’s *t*-tests; ns: *p* > 0.05; *: *p* < 0.05; ** *p* < 0.01.

Oscillatory potentials (OPs) are low amplitude, high frequency waves superimposed over the rising phase of the ERG b-wave and observed after bright stimuli. Previous studies indicate that OPs originate from rhythmic activity in the IPL downstream of the bipolar cells (Ogden, 1973). Because the light response of a dark-adapted retina is primarily rod-driven, the OPs are indirectly correlated to RBC output. Therefore, we normalized the OP amplitude by the b-wave amplitude (OP/b-wave ratio) of the corresponding ERG recording. Normalized TPBG-KO OPs were reduced compared to WT (**Figure 2D and E**). TPBG is primarily expressed in RBCs, therefore an effect of TPBG-KO on the OPs is most likely to be due to reduced RBC output onto downstream neurons, though an effect mediated by the TPBG-expressing amacrine cells cannot be ruled out.

### TPBG is not required for mGluR6-driven responses in RBC dendrites

We examined the expression and localization patterns of key pre- and post-synaptic proteins that are required for synaptic transmission between rod photoreceptors and RBCs. RBCs receive synaptic input from rod photoreceptors via a sign-inverting signal transduction cascade localized to the tips of RBC dendrites. Glutamate released from rod photoreceptors is detected by the group III metabotropic glutamate receptor mGluR6 (Masu et al., 1995; Nakajima et al., 1993; Vardi, Duvoisin, Wu, & Sterling, 2000; Xu et al., 2012), that is negatively coupled to the TRPM1 cation channel (Koike, Numata, Ueda, Mori, & Furukawa, 2010; Koike, Obara, et al., 2010; Morgans et al., 2009; Shen et al., 2009). Thus, a light stimulus relieves TRPM1 tonic inhibition, permitting cation influx and subsequent RBC depolarization.

First, we looked at the spatial relationship between presynaptic elements in the rod spherule and the tips of the RBC dendrites (**Figure 3A**) using an antibody against the rod L-type Ca^2+^ channel subunit Ca_v_1.4 to label the horseshoe-shaped active zone and an antibody that labels GPR179 in RBC dendritic tips (Hasan, Ray, & Gregg, 2016). In normal rod-RBC synapses, GPR179 labeling appears as puncta within the concavity of the horseshoe-shaped Ca_v_1.4 immunofluorescence. We quantified the spatial relationship between rod active zones and RBC dendritic tips by measuring the distance between the peak intensity for each fluorescent channel (**Figure 3B**). We found no difference between WT (blue; *n* = 66 synapses from eight animals) and TPBG-KO (KO; orange; *n* = 54 synapses from seven animals) synaptic distance (**Figure 3C**; WT: 0.43 ± 0.006 μm; KO: 0.43 ± 0.004 μm; Welch’s *t*-test: *p* = 0.96, *t*(117.60) = 0.052), suggesting that RBC dendrites form morphologically normal invaginations into rod spherules in the TPBG-KO mouse.

**Figure 3:**
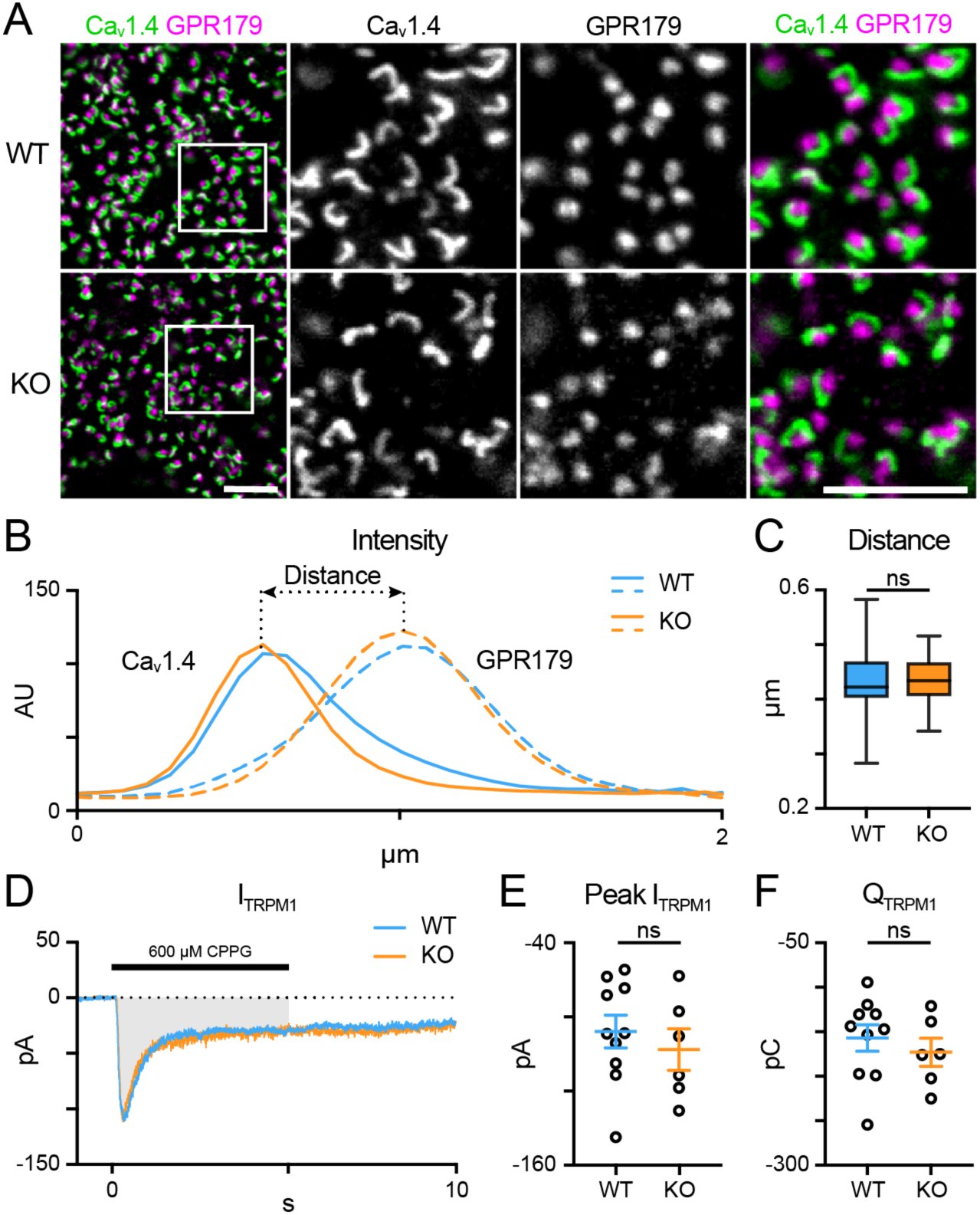
TPBG is not required for mGluR6-driven responses in RBC dendrites. (**A**) Immunofluorescence confocal microscopy from WT and TPBG-KO whole mount retinas using antibodies against the rod presynaptic calcium channel Ca_v_1.4 (green) and the RBC post-synaptic protein GPR179 (magenta). Square boxes in the leftmost images indicate regions of interest expanded in the other images. Z-projections of two sections were used with a Z-step distance of 0.297 μm for a total thickness of 0.594 μm. Scale bar is 5 μm. **(B)** Plot of fluorescence intensity over distance from Ca_v_1.4 (left) and GPR179 (right) channels from WT (blue; *n* = 66 synapses) and TPBG-KO (KO; orange; *n* = 54 synapses). **(C)** Quantification of synaptic distance. Box plot extends from the first to the third quartile and the whiskers extend across all points. The line in the box represents the median. **(D-F)** Chemically simulated light responses were recorded in WT (blue; *n* = 10) and TPBG-KO (KO; orange; *n* = 6) RBCs following a pulse of 600 µM CPPG for 5 seconds. **(D)** Example CPPG-induced TRPM1 currents. The gray bar represents the region across which charge transfer was calculated. **(E)** Quantification of peak CPPG-induced TRPM1 current (I_TRPM1_) and **(F)** charge transfer (Q_TRPM1_) over the five-second duration of the CPPG pulse. For **E** and **F**, open circles represent single cells and colored bars represent the mean ± SEM. Statistical significance was determined using Welch’s *t*-tests; ns: *p* > 0.05.

Scotopic ERG recordings revealed increased b-waves in TPBG-KO mice (**Figure 2**) indicating an enhanced light response in KO RBCs. Because the RBC light response is initiated by TRPM1, we compared TRPM1 currents between WT (blue; *n* = 10) and TPBG-KO (KO; orange; *n* = 6) RBCs by recording chemically simulated light responses in retinal slices in whole-cell voltage-clamp (**Figure 3D-F**). In this technique, bath application of the mGluR6 agonist L-AP4 maintains the retina in a simulated dark-adapted state, and brief puffs of the mGluR6 antagonist, CPPG, onto the dendrites simulates a light flash and evokes TRPM1 currents (**Figure 3D**). TPBG-KO did not alter the CPPG-induced TRPM1 peak current (**Figure 3E**; WT: -88.24 ± 8.92 pA; KO: -98.04 ± 11.28 pA; Welch’s *t*-test: *p* = 0.51, *t*(10.85) = 0.68) or charge transfer (**Figure 3F**; WT: -157.10 ± 47.01 pC; KO: -173.10 ± 38.62 pC; Welch’s *t*-test: *p* = 0.47, *t*(12.40) = 0.74). These results suggest that TPBG is not required for the formation of functional rod-RBC synapses and that the larger scotopic ERG b-waves do not reflect enhanced TRPM1 currents in the TPBG-KO.

### Synaptic ribbons are smaller in TPBG-KO RBCs compared to WT

Oscillatory potentials are thought to be generated by the inner retina, and RBCs drive the majority of downstream activity under scotopic conditions; thus, the attenuation of scotopic OPs in the TPBG-KO suggests a deficit in RBC output to third-order neurons. Like rods, RBCs signal to downstream cells using ribbon synapses. A depolarized potential entering the axon terminal prompts the opening of L-type voltage-gated Ca^2+^ channels clustered beneath the ribbons and the resulting influx of Ca^2+^ triggers the fusion (exocytosis) of ribbon-tethered synaptic vesicles (Gersdorff, Vardi, Matthews, & Sterling, 1996). This process requires a close spatial relationship between the Ca^2+^ channels the synaptic ribbons (Jarsky, Tian, & Singer, 2010). We used triple immunofluorescence microscopy to examine the relative localization and expression patterns of the L-type Ca^2+^ channel subunit, Ca_v_1.4, and the primary structural synaptic ribbon protein RIBEYE (**Figure 4A-C**). Because neither Ca_v_1.4 nor RIBEYE are specific to RBC axon terminals, we isolated RBC-specific immunofluorescence using a binary mask derived from PKCα immunofluorescence to exclude labeling from other bipolar cells in the IPL. Ca_v_1.4 and RIBEYE show punctate immunofluorescence in RBC axon terminals and are co-localized or closely apposed (**Figure 4A**). Mander’s coefficients (M1 and M2) were calculated to examine colocalization between Ca_v_1.4 and RIBEYE puncta in both control (PKCα-negative) terminals and RBC (PKCα-positive) terminals (**Figure 4B**). We found no differences in co-localization coefficients between WT and TPBG-KO control or RBC axon terminals: WT (blue; *n* = 134 control puncta and 100 RBC puncta) and TPBG-KO (orange; *n* = 327 control puncta and 119 RBC puncta), control M1 (WT: 0.53 ± 0.022 arbitrary units (AU); KO: 0.52 ± 0.013 AU; Welch’s *t*-test: *p* = 0.60, *t*(237.2) = 0.53), control M2 (WT: 0.40 ± 0.015 AU; KO: 0.42 ± 0.011 AU; Welch’s *t*-test: *p* = 0.28, *t*(288.1) = 1.09), RBC M1 (WT: 0.66 ± 0.016 AU; KO: 0.64 ± 0.015 AU; Welch’s *t*-test: *p* = 0.47, *t*(208.3) = 0.73), RBC M2 (WT: 0.32 ± 0.01 AU; KO: 0.31 ± 0.01 AU; Welch’s *t*-test: *p* = 0.76, *t*(216.9) = 0.31). Additionally, we found that the number of ribeye puncta per unit area was unchanged in WT (blue; *n* = 90 axon terminals) and TPBG-KO (KO; orange; *n* = 109 axon terminals) RBC terminals (**Figure 4C**; WT: 2.07 ± 0.66 puncta/μm^2^; KO: 2.03 ± 0.58 puncta/μm^2^; Welch’s *t*-test: *p* = 0.70, *t*(177.8) = 3.92). We also compared synaptotagmin I/II immunofluorescence between RBC axon terminals in WT and TPBG-KO retinas and saw no difference (not shown; WT: 1.352 ± 0.0486 AU; KO: 1.446 ± 0.0560; Welch’s t-test: p-value = 0.209, t(52.63) = 1.271).

**Figure 4:**
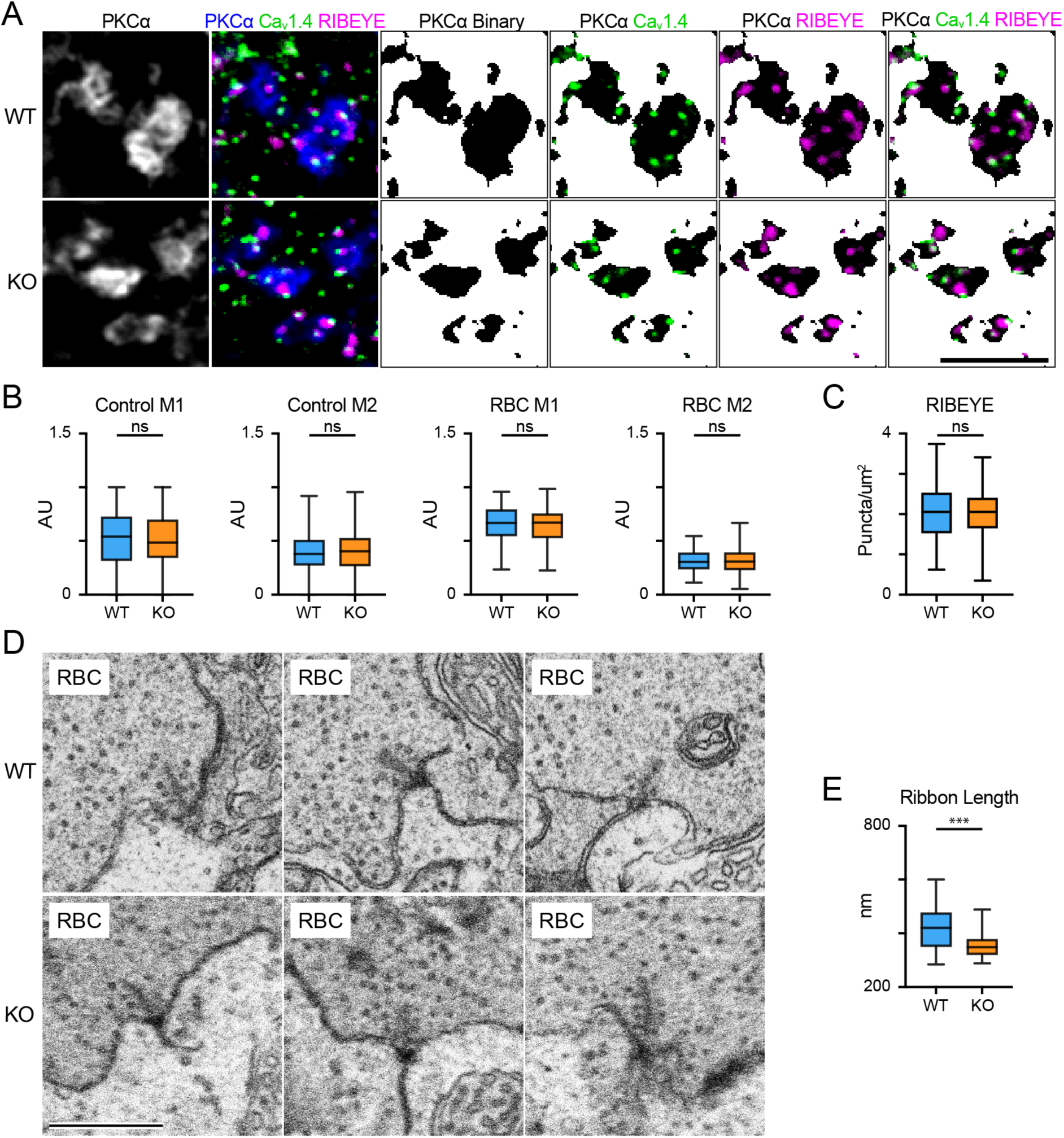
RBC synaptic ribbons are smaller in the TPBG-KO retina. (**A**) Triple immunofluorescence confocal microscopy of WT and TPBG-KO (KO) whole mount retinas using antibodies against PKCα (blue), Ca_v_1.4 (Green) and RIBEYE (magenta). To isolate labeling in RBC axon terminals, binary images were created using the PKCα channel and overlayed over the other two channels to generate binary masks. Z-projections of two sections were used with a Z-step distance of 0.297 μm for a total thickness of 0.547 μm. The scale bar is 5 μm. (**B**) Quantification of Ca_v_1.4 and RIBEYE colocalization in WT (blue) and TPBG-KO (KO; orange) using M1 and M2 Mander’s coefficients in control (WT: *n* = 134 puncta; KO: *n* = 327 puncta) and RBC axon terminals (WT: *n* = 100 puncta; KO: *n* = 119 puncta). (**C**) Quantification of RIBEYE puncta counts in WT (blue; *n* = 90 terminals) and TPBG-KO (KO; orange; *n* = 109 terminals). (D) Example FIB-SEM images showing RBC active zones from WT (top) and TPBG-KO (KO; bottom) RBC axon terminals. Scale bar is 500 nm. (**E**) Quantification of synaptic ribbon length in WT (blue; *n* = 33 ribbons) and TPBG-KO (KO; orange; *n* = 38 ribbons) measured from the membrane. Box plots extend from the first to the third quartile and the whiskers extend across all points. The line in the box represents the median.

To examine synaptic ribbons in RBC axon terminals, serial electron micrographs were collected from a volume each of WT and TPBG-KO retina using a focused ion beam scanning electron microscope (FIB-SEM). Bipolar cell terminals closest to the ganglion cell layer were identified as putative RBC axon terminals. The general ultrastructure of the RBC axon terminals, including the synaptic vesicles, synaptic ribbons, and postsynaptic elements was similar in WT and TPBG-KO retina. To estimate synaptic ribbon size, we measured the maximum distance that each ribbon extended into the cytoplasm from its attachment site on the plasma membrane in WT (blue; *n* = 33 ribbons) and TPBG-KO (KO; orange; *n* = 38 ribbons) and found that synaptic ribbons were 15% smaller in TPBG-KO RBCs compared to WT RBCs (**Figure 4E**; WT: 416.8 ± 13.78 nm; KO: 355.7 ± 8.85 nm; Welch’s *t*-test: *p* = 0.0004, *t*(55.65) = 3.74).

### TPBG is required for efficient exocytosis in RBC synaptic terminals

To examine the effect of TPBG-KO on RBC synaptic output, we recorded time-resolved membrane capacitance (C_m_) before and after a depolarizing pulse using the “sine + DC” method (Gillis, 2000; Oltedal & Hartveit, 2010). High-resolution membrane capacitance changes (ΔC_m_) after a depolarizing pulse strongly correlate with vesicle exocytosis in neuroendocrine cells (Klyachko & Jackson, 2002; Neher & Marty, 1982) and glutamate release from bipolar cell axon terminals (Kim & von Gersdorff, 2016). A sinusoidal voltage command (200 Hz at 30 mV trough-to-peak) was superimposed over the holding voltage of -80 mV and exocytosis was evoked by depolarizing the cell to -10 mV to open the voltage-gated Ca^2+^ channels (**Figure 5: Supplementary Figure A)**. Membrane capacitance (C_m_), membrane resistance (R_m_), and series resistance (R_s_) were recorded before and after the pulse and ΔC_m_ was measured as the change in C_m_ after exocytosis. Changes in R_m_ and R_s_ were not correlated with changes in C_m_ (**Figure 5: Supplementary Figure B**).

**Figure 5:**
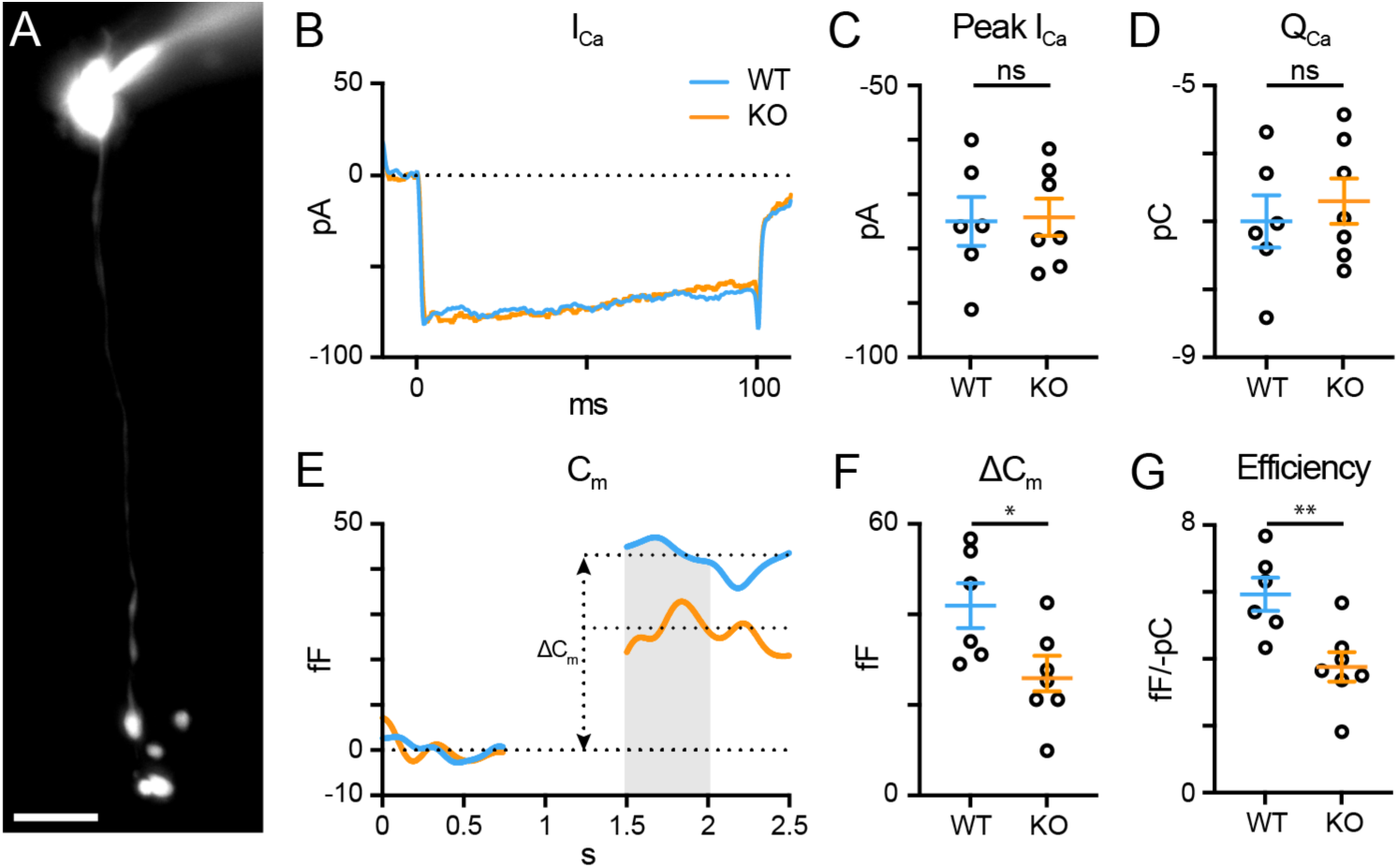
TPBG is required for efficient synaptic vesicle release in RBC axon terminals. Whole-cell voltage-clamp recordings were performed in WT (blue; *n* = 6) and TPBG-KO (KO; orange; *n* = 7) RBCs. **(A)** Epifluorescence micrograph of an RBC filled with Alexa Fluor 594 Hydrazide after patch-clamp recording. RBCs were patched and cell type was verified *post-hoc* using epifluorescence microscopy. The scale bar is 10 μm. **(B)** Average voltage-gated Ca^2+^ current traces recorded after a depolarizing pulse from -80 mV to -10 mV for 100 ms. **(C)** Quantification of peak voltage-gated Ca^2+^ current amplitudes (I_Ca_) and **(D)** charge transfer (Q_Ca_) over the duration of the pulse. **(E)** Average membrane capacitance (C_m_) recordings using a 200 Hz sine + DC protocol. ΔC_m_ was measured as the difference between baselined resting C_m_ before and after the depolarizing pulse. Data immediately after the pulse was excluded until C_m_ stabilized. The gray bar represents the region across which C_m_ values were averaged. **(F)** Quantification of ΔC_m_. **(G)** Quantification of exocytosis efficiency calculated as the absolute value of the change in C_m_ per unit Q_Ca_. Ames medium was supplemented with 1 mM CaCl_2_ and (in µM): 0.5 strychnine, 3 gabazine, 50 TPMPA, and 4 L-AP4. Open circles in bar graphs represent single cells and bars represent the mean ± SEM. Statistical significance was determined using Welch’s *t*-tests; ns: *p* > 0.05; *: *p* < 0.05; **: *p* < 0.01.

C_m_ measurements in RBCs are complicated by the attenuation and filtering of the sinusoidal voltage command amplitude down their narrow axons (Oltedal & Hartveit, 2010; Oltedal, Veruki, & Hartveit, 2009). We sought to confirm that a low-frequency sinusoidal command reduces attenuation and maximizes RBC C_m_ changes compared to higher frequencies (**Figure 5: Supplementary Figure C-F**) using 200 (n = 6), 400 (n = 4), 800 (n = 4), and 1600 (n = 4) Hz commands. We found no difference in the voltage-gated Ca^2+^ current amplitude (I_Ca_) or charge transfer (Q_Ca_) across frequencies (not shown), but the ΔC_m_ (**Figure 5: Supplementary Figure C-D**; 200 Hz: 41.94 ± 4.95 fF; 400 Hz: 19.13 ± 4.06 fF; 800 Hz: 4.05 ± 2.91 fF; 1600 Hz: -4.23 ± 1.40; Brown-Forsythe ANOVA: p < 0.0001, 34.12(2.00, 10.87)) and the resting C_m_ (**Figure 5: Supplementary Figure E-F**; 200 Hz: 5.97 ± 0.23 pF, 400 Hz: 5.73 ± 0.12 pF, 800 Hz: 4.81 ± 0.15 pF, 1600 Hz: 4.04 ± 0.12 pF; Brown-Forsythe ANOVA: p < 0.0001, 25.06(3.00, 11.34)) were greater at 200 Hz compared to higher frequencies. These supplementary results support our decision to examine exocytosis in RBCs using a 200 Hz “sine + DC” protocol.

To examine RBC synaptic vesicle exocytosis, we used whole-cell voltage-clamp to compare WT (blue; n = 6) and TPBG-KO (KO; orange; n = 7) voltage-gated Ca^2+^ currents and subsequent changes in membrane capacitance (**Figure 5**). Cells where C_m_, R_m_, and R_s_ were highly correlated were discarded. The synaptic blockers L-AP4, strychnine, TPMPA, and gabazine were used to block spontaneous activity and help increase and stabilize R_m_ during recordings. The passive properties of R_m_ (WT: 1.603 ± 0.24 GΩ; KO: 1.630 ± 0.16 GΩ; Welch’s *t*-test: p = 0.93, *t*(8.84) = 0.097) and R_s_ (WT: 50.54 ± 2.33 MΩ; KO: 49.61 ± 1.91 MΩ; Welch’s *t*-test: p = 0.76, *t*(10.16) = 0.31) were not different between WT and TPBG-KO RBCs. As long as R_m_ is significantly greater than R_s_, ΔC_m_ should be a faithful approximation of overall exocytosis (Gillis, 2000).

TPBG-KO had no effect on the voltage-gated Ca^2+^ currents (**Figure 5B-C**; WT: -74.99 ± 4.49 pA, KO: -74.24 ± 3.41 pA; Welch’s *t*-test: *p* = 0.90, *t*(9.73) = 0.13) or charge transfer (**Figure 5D**; WT: -7.0 ± 0.38 pC, KO: -6.7 ± 0.33 pC; Welch’s *t*-test: *p* = 0.57, *t*(10.43) = 0.58). However, TPBG-KO resulted in a reduction in ΔC_m_ compared to WT (**Figure 5E-F**; WT: 41.94 ± 4.95 fF, KO: 25.90 ± 3.91 fF; Welch’s *t*-test: *p* = 0.029, *t*(9.95) = 2.55), suggesting a decrease in synaptic vesicle exocytosis in the KO. Most interestingly, the efficiency of exocytosis (the ratio of ΔC_m_ per unit Q_Ca_) is also reduced in the KO (**Figure 5G**; WT: 5.93 ± 0.50 fF/-pC, KO: 3.76 ± 0.44 fF/-pC; Welch’s *t*-test: *p* = 0.0077, *t*(10.52) = 3.28), indicating that TPBG-KO alters the efficiency of RBC vesicle exocytosis without changing Ca^2+^ influx.

### TPBG does not alter RBC input resistance or excitability

TPBG-KO amplifies the RBC light response and reduces synaptic vesicle exocytosis, suggesting that TPBG-KO may reduce the gain of RBC synaptic release by shifting RBC excitability. To compare the excitability between WT (blue; *n* = 9) and TPBG-KO (KO; orange; *n* = 11) RBCs, we used current injections (from -80 to 80 pA in 20 pA intervals) in whole-cell current-clamp mode to construct voltage-current relationships and calculate the peak and sustained input resistances (R_in_). For these experiments, we used the synaptic blockers L-AP4, strychnine, TPMPA, and gabazine to isolate RBC-specific voltage responses. **Figure 6A** shows voltage responses after representative current injections of -80 pA, 0 pA, and +80 pA. The resting membrane potential (V_m_) was unchanged in the KO (**Figure 6B**; WT: -58.02 ± 4.91 mV, KO: -56.35 ± 4.66 mV; Welch’s *t*-test: *p* = 0.81, *t*(17.53) = 0.25). We found no difference in peak (**Figure 6C-D**; WT: 1.40 ± 0.13 GΩ, KO: 1.39 ± 0.09 GΩ; Welch’s *t*-test: *p* = 0.97, *t*(14.94) = 0.04) or sustained R_in_ (**Figure 6E-F**; WT: 1.13 ± 0.11 GΩ, KO: 1.05 ± 0.07 GΩ; Welch’s *t*-test: *p* = 0.58, *t*(14.28) = 0.56) between the WT and KO RBCs. These results indicate that a shift in R_in_ and RBC excitability in the TPBG-KO do not explain the reduced exocytosis.

**Figure 6:**
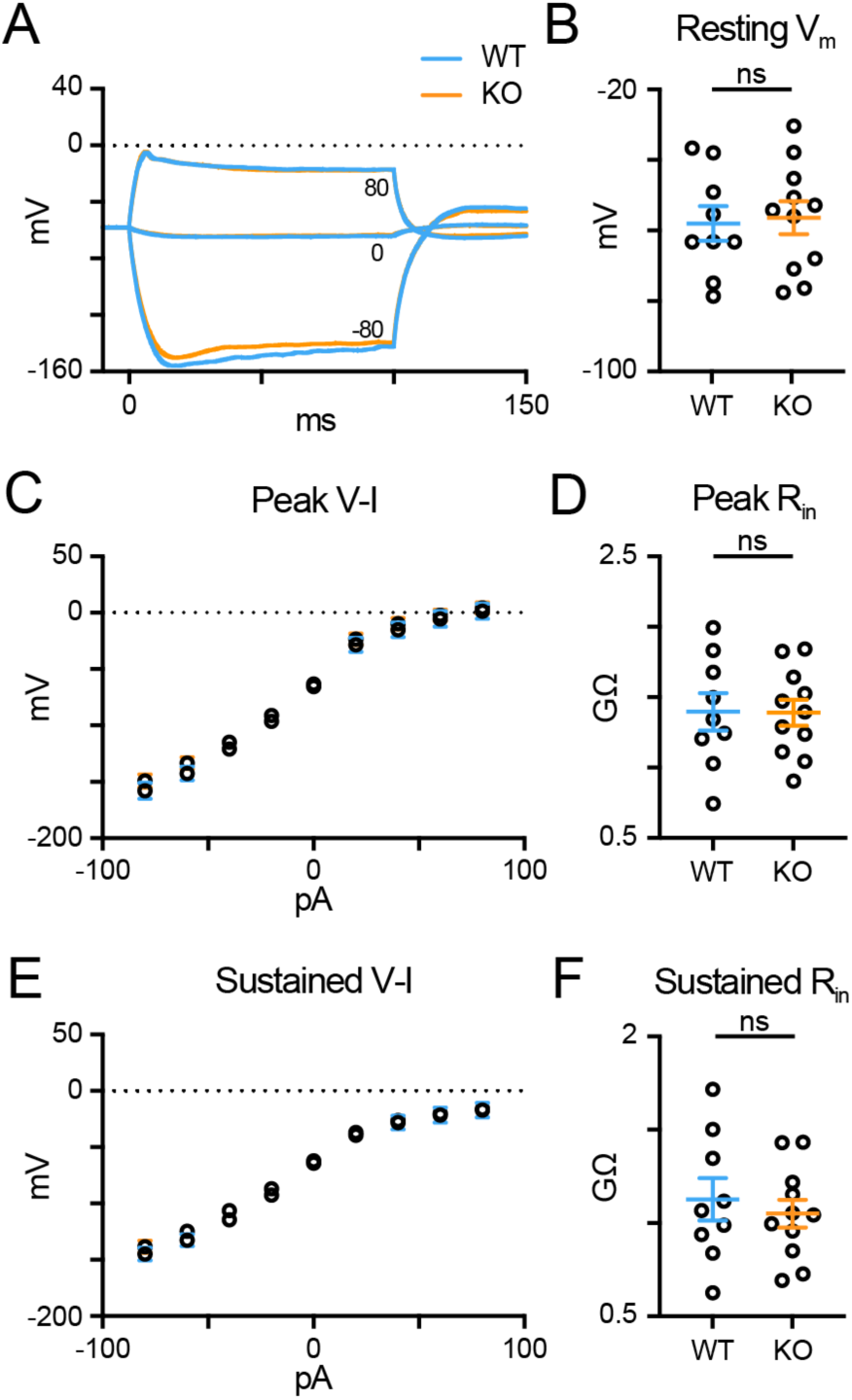
TPBG does not alter RBC input resistance or excitability. Whole-cell current-clamp recordings in WT (blue; *n* = 9) and TPBG-KO (KO; orange; *n* = 11) RBCs were used to construct a current-voltage relationship and calculate input resistance (R_in_). A current injection protocol from -80 pA to 80 pA in 20 pA steps was used with a K-gluconate-based internal solution. **(A)** Average voltage traces following -80 pA, 0 pA, and 80 pA current injections for 100 ms. **(B)** Resting membrane voltages recorded immediately after entering current clamp mode. **(C)** A plot of the voltage-current (V-I) relationship measured at the peak voltage during current injections. **(D)** Quantification of peak R_in_. (**E**) A plot of the sustained V-I relationship measured as the average voltage calculated over the last 10 ms of the current pulse. **(F)** Quantification of the sustained R_in_. Peak and sustained R_in_ values were calculated by measuring the slope of the linear portion of the V-I curves (from -40 to 40 pA). Ames medium was supplemented with 1 mM CaCl_2_ and (in µM): 0.5 strychnine, 3 gabazine, 50 TPMPA, and 4 L-AP4. For V-I curves, open circles represent the mean and error bars represent SEM. For quantification, open circles represent single cells and bars represent the mean ± SEM. Statistical significance was determined using Welch’s *t*-tests; ns: *p* > 0.05.

### TPBG-KO reduces reciprocal inhibitory feedback

RBCs drive retinal output via glutamate release into the IPL, which stimulates Ca^2+^-permeable AMPA receptors in downstream AII amacrine cells (AII-ACs). RBC glutamate release is also detected by A17 amacrine cells (A17-ACs), which when depolarized, release GABA back onto RBC synaptic terminals where it is sensed by GABA_A_ and GABA_A_ρ (also called GABA_C_) Cl^-^ channels (**Figure 7A**). An influx of Cl^-^ into the axon terminal through open GABA channels provides inhibitory feedback by hyperpolarizing the RBC and shaping further RBC glutamate release (Chávez, Grimes, & Diamond, 2010; Chávez, Singer, & Diamond, 2006; Euler & Masland, 2000; E. Hartveit, 1996; Espen Hartveit, 1999; Singer & Diamond, 2003). Each RBC active zone forms a dyad synapse with one AII-AC and one A17-AC (Famiglietti & Kolb, 1975; Grimes, Zhang, Graydon, Kachar, & Diamond, 2010; McGuire, Stevens, & Sterling, 1984; Raviola & Dacheux, 1987). As A17-mediated reciprocal feedback is driven by RBC glutamate release, the inhibitory feedback currents in the RBCs are an indirect measure of glutamate release from RBCs and would be expected to be reduced concomitantly with reduced RBC exocytosis in the TPBG-KO.

**Figure 7:**
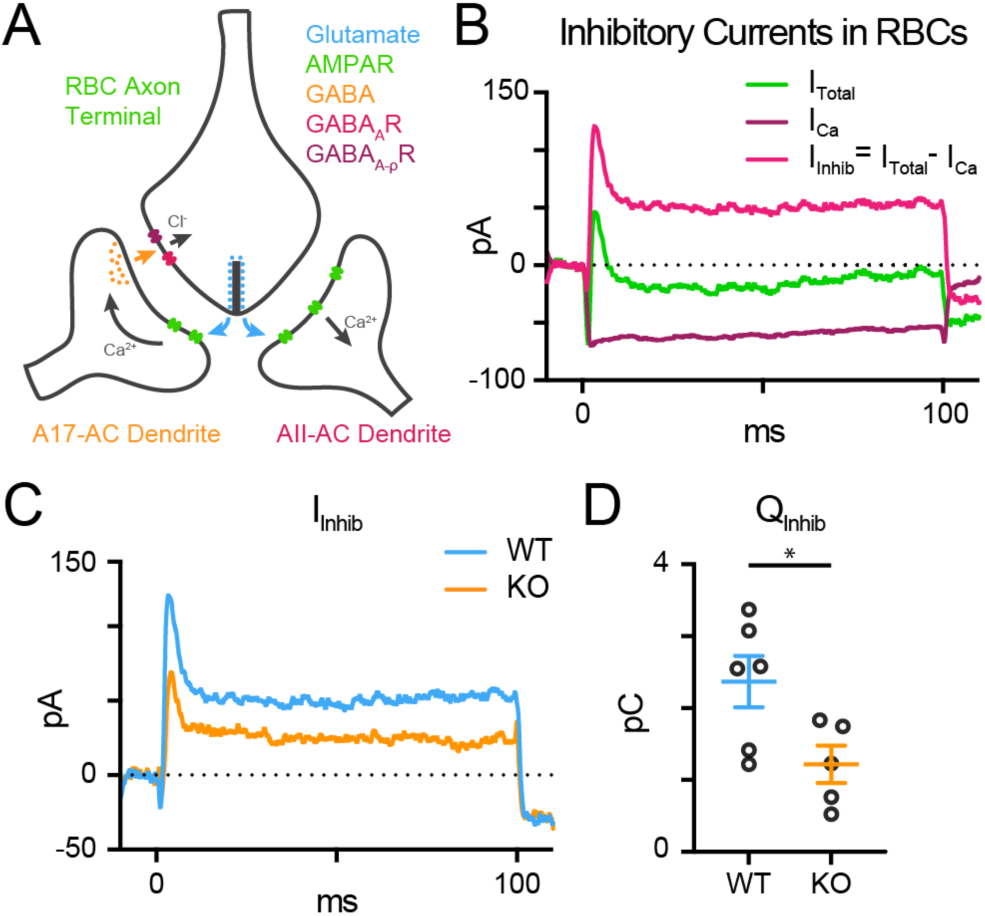
TPBG-KO reduces reciprocal inhibitory feedback. **(A)** The dyad synapse arrangement between one RBC, one AII-AC, and one A17-AC in the IPL. Glutamate release by RBCs is detected by Ca^2+^-permeable AMPA receptors (AMPARs) in AII-AC and A17-AC dendrites. A17-ACs then release GABA back onto the synaptic cleft which is sensed by GABA_A_ receptors (GABA_A_Rs) and GABA_A_ρ receptors (GABA_A_ρRs) in the RBC axon terminals. **(B)** Inhibitory currents are superimposed over voltage-gated Ca^2+^ currents. Reciprocal feedback in RBCs was evoked with a 100 ms depolarizing pulse from -80 mV to -10 mV. In the absence of GABA receptor blockers, this depolarizing pulse elicits an inward voltage-gated Ca^2+^ current that is quickly subsumed by outward inhibitory currents (I_Total_; green). These outward currents are abolished by the application of the GABA receptor blockers gabazine and TPMPA (I_Ca_; magenta). Inhibitory currents (I_Inhib_; magenta) can be isolated by subtracting I_Ca_ (maroon) from I_Total_ (green). **(C)** Average whole-cell patch-clamp recordings from WT (blue; *n* = 6) and TPBG-KO (KO; orange; *n* = 5) RBCs to compare reciprocal inhibitory feedback (I_Inhib_ = I_Total_ – I_Ca_) after a depolarizing pulse from -80 mV to -10 mV for 100 ms. The gray bar represents the region across which the charge transfer was calculated. **(D)** Quantification of the charge transfer (Q_Inhib_) measured from baseline over the duration of the 100 ms stimulus. Ames medium was supplemented with 1 mM CaCl_2_, 0.5 µM strychnine and 4 µM L-AP4. Open circles in bar graphs represent single cells and bars represent the mean ± SEM. Statistical significance was determined using Welch’s *t*-tests; *: *p* < 0.05.

To test the hypothesis that reduced RBC glutamate release results in smaller inhibitory feedback currents in the TPBG-KO, we used whole-cell voltage-clamp to examine the reciprocal inhibitory response to a depolarizing pulse from -80 mV to -10 mV for 100 ms in WT (blue; *n* = 6) and TPBG-KO (KO; orange; *n* = 5) RBCs. L-AP4 was used to block spontaneous excitatory inputs and strychnine was used to block glycinergic inhibition. In the absence of GABA receptor blockers, outward currents can be seen superimposed on the evoked voltage-gated Ca^2+^ current (**Figure 7B; Green;** I_Total_). Inhibitory currents (magenta; I_Inhib_) can be isolated by subtracting voltage-gated Ca^2+^ currents (maroon; I_Ca_)from the total current. The charge transfer across the duration of the stimulus was reduced in the TPBG-KO compared to WT (**Figure 7D**; WT: 2.35 ± 0.37 pC; KO: 1.13 ± 0.22 pC; Welch’s *t*-test: *p* = 0.022, *t*(7.83) = 2.83). This supports our previous result that the TPBG-KO suppresses RBC vesicle exocytosis and glutamate release (see **Figure 5F**).

It is possible that reciprocal feedback is altered in TPBG-KO due to disrupted expression or clustering of GABA receptors in RBC axon terminals. RBCs express both GABA_A_α1 and GABA_A_ρ1 receptor subunits in their axon terminals (Fletcher, Koulen, & Wässle, 1998; Palmer, 2006). To see whether TPBG-KO alters GABA receptor expression patterns, we used double immunofluorescence labeling of WT and TPBG-KO whole mount retinas using an antibody against PKCα and antibodies against GABA_A_α1 (WT: *n* = 18 terminals from three animals; KO *n* = 28 terminals from four animals) and GABA_A_ρ1 (WT: *n* = 20 terminals from three animals; KO *n* = 27 terminals from three animals) receptor subunits (**Figure 8**). As previously, we isolated RBC axon terminal-specific labeling using a binary mask derived from PKCα immunofluorescence. Next, we counted the GABA puncta per unit area in PKCα-positive axon terminals. We did not observe any major differences in puncta/μm^2^ between WT and TPBG-KO (KO) retinas labeled against either GABA_A_α1 (**Figure 8A-B**; WT: 1.21 ± 0.05 puncta/μm^2^; KO: 1.40 ± 0.09 puncta/μm^2^; Welch’s *t*-test: *p* = 0.064, *t*(39.73) = 1.91) or GABA_A_ρ1 (**Figure 8C-D**; WT: 1.03 ± 0.03 puncta/μm^2^; KO: 0.98 ± 0.03 puncta/μm^2^; Welch’s *t*-test: *p* = 0.20, *t*(41.11) = 1.29), indicating that impaired GABA receptor expression is likely not reducing inhibitory feedback. Taken together, our data supports the conclusion that TPBG is required for normal synaptic ribbon development and efficient glutamate release from RBC axon terminals.

**Figure 8:**
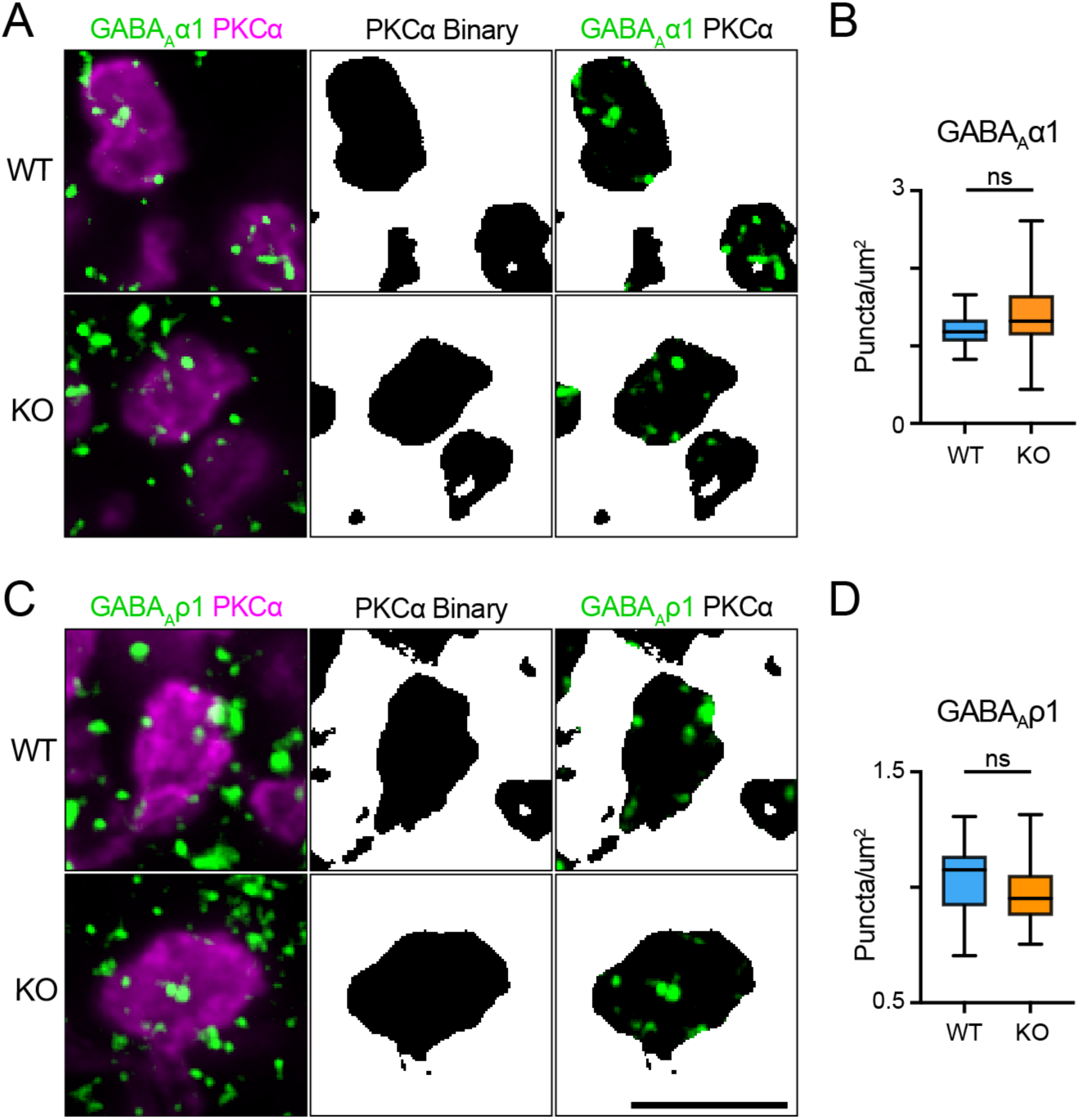
TPBG is not required for localization of GABA receptor immunofluorescence to RBC axon terminals. Immunofluorescence confocal microscopy of WT and TPBG-KO (KO) whole mount retinas using an antibody against PKCα (magenta) and antibodies against either the GABA_A_α1 subunit (**A**; green; WT: *n* = 18 terminals; KO *n* = 28 terminals) or the GABA_A_ρ1 subunit (**C**; green; WT: *n* = 20 terminals; KO *n* = 27 terminals). To isolate labeling in RBC axon terminals, binaries were created using the PKCα channel and overlayed over the other channels to generate binary masks. For all images, Z-projections of two sections were used with a Z-step distance of 0.297 μm for a total thickness of 0.547 μm. Scale bar is 5 μm. **(B)** Quantification of GABA_A_α1 subunit puncta/μm^2^. **(D)** Quantification of GABA_A_α1 subunit puncta/μm^2^. Box plots extend from the first to the third quartile and the whiskers extend across all points. The line in the box represents the median. Statistical significance was determined using Welch’s *t*-tests; ns: *p* > 0.05.

## Discussion

In this study, we characterized the functional consequences of genetic knockout of TPBG in the mouse retina using immunofluorescence and electrophysiological approaches. TPBG-KO did not alter gross RBC morphology in the OPL or IPL as PKCα-labeled RBC dendrites and axon terminals appear normal in both conditions. Because TPBG is localized to both the dendrites and axon terminals of RBCs (Wakeham et al., 2020), we sought to examine the effect of TPBG-KO on the RBC synaptic structure and physiology in both compartments. Though ERG b-waves were larger in the TPBG-KO, dendritic TRPM1 currents activated by CPPG did not change. In contrast to the larger b-waves, the scotopic OPs were smaller in the KO, suggesting changes in RBC ribbon synapse function. Accordingly, time-resolved C_m_ measurements and whole-cell recording indicated that synaptic vesicle exocytosis and GABAergic feedback are reduced at TPBG-KO RBC synapses, and synaptic ribbons are shorter. Importantly, we did not detect differences in presynaptic voltage-gated Ca^2+^ current amplitude or changes in Ca^2+^ channel, GABA receptor, or RIBEYE expression and localization. We propose that TPBG plays an important role in fine tuning the synaptic architecture of RBC ribbon synapses to increase exocytosis efficiency.

### TPBG and RBC dendrites

We found that TPBG is not required for the normal insertion of RBC dendritic tips inside rod spherules or for normal depolarizing currents through TRPM1 cation channels. Our immunofluorescence and electrophysiological examination of RBC dendrites suggest that TPBG is not essential for basic synaptic function in that compartment. That being said, TPBG immunofluorescence reveals that large amounts of TPBG are present in RBC dendrites, suggesting some role in dendritic development or physiology.

TPBG regulates dendritic arborization in mouse olfactory bulb granule cells (Yoshihara et al., 2012). In these neurons, TPBG-KO reduced dendritic branching and dendritic migration to synaptic targets. Lentiviral overexpression of TPBG increased dendritic branching complexity and restored dendritic arborization deficits induced by nasal occlusion. In the retina, TPBG may also regulate RBC dendritic branching complexity in the OPL. TPBG is a known regulator of Wnt signaling (He et al., 2015; Kagermeier-Schenk et al., 2011), which is required for functional synaptic targeting and OPL lamination in RBCs, therefore TPBG may promote RBC dendritic development through control of the Wnt pathway (Tsuboi, 2020). We didn’t detect a structural or physiological phenotype in RBC dendrites, but thorough evaluation of RBC dendritic maturation and branching using sparse labeling techniques might reveal a more subtle effect of TPBG-KO on dendritic architecture.

### TPBG and RBC synaptic vesicle exocytosis

We found a reduction in the OPs isolated from scotopic ERG recordings in TPBG-KO mice, which we interpret as evidence for decreased RBC output to third-order neurons. While we cannot rule out a contribution of the TPBG-expressing amacrine cells to the reduced OP amplitudes in the TPBG-KO, the similarity of the OP phenotype to that seen in the PKCα-KO (Xiong et al., 2015) suggests that it is due to an RBC defect. Furthermore, pharmacological dissection of ERG components points to the RBC to AII/A17 reciprocal synapse as the origin of the OPs (Liao, Liu, Milla-Navarro, Villa, & Germain, 2023; Wachtmeister, 1998). Early studies implicated interactions between bipolar cell axon terminals, amacrine cell processes, and ganglion cell dendrites as potential generators of the OPs (Ogden, 1973). Intravitreal injections of glycine into rabbit retinas resulted in morphological changes in many ACs and disappearance of the OPs from the ERG (Korol, Leuenberger, Englert, & Babel, 1975) while application of neurotoxic kainate to ganglion cell dendrites diminishes OP amplitude (Vaegan & Millar, 1994). A more recent study unequivocally implicates multiple distinct generators in the production of early, intermediate, and late OP peaks (Dong, Agey, & Hare, 2004). The use of APB and CNQX to block synaptic inputs to ON and OFF-bipolar cells significantly diminished intermediate and later OPs but left early OPs intact, suggesting that early OPs are generated by photoreceptors. Intermediate and late OPs have differential sensitivity to tetrodotoxin, which abolished only late OPs, indicating that intermediate OPs are generated by action-potential-independent interactions and late OPs are generated by action-potential-dependent ones (Dong et al., 2004). Our OP data shows a much larger attenuation of TPBG-KO OP amplitude in intermediate and late OPs compared to early ones. Thus, the results of these studies support our conclusion that reduced scotopic OPs are likely due to suppressed rhythmic activity of third-order neurons due to reduced RBC glutamate release.

By simultaneously recording the presynaptic voltage-gated Ca^2+^ currents and subsequent membrane capacitance changes evoked by RBC depolarization, we showed that reduced exocytosis in TPBG-KO RBCs compared to WT, despite no change in presynaptic Ca^2+^ currents, indicating reduced exocytosis efficiency in the TPBG-KO. The Ca^2+^ current amplitudes we recorded (-75 pA) following a depolarizing step are significantly larger than those previously reported (-30 pA by (Protti & Llano, 1998) and -20 pA by (Singer & Diamond, 2003)). We also recorded larger C_m_ jumps (42 fF in WT) than those reported from dissociated RBCs (30 fF in (Zhou, Wan, Thakur, & Heidelberger, 2006)). We believe these differences can be attributed to higher recording temperatures (33° C vs 21° C) and our use of older animals. With a mean resting C_m_ of 5.97 pF and a mean R_s_ of 50.04 MΩ, we can calculate an average voltage-clamp speed of 298.74 μs – sufficiently fast to clamp the membrane to a 200 Hz sine wave with a 5 ms period. The reduced exocytosis in the TPBG-KO, despite the same Ca^2+^ currents as in WT, suggests that calcium influx is not as tightly coupled to the synaptic vesicle exocytosis in the TPBG-KO, resulting in reduced exocytosis efficiency.

Exocytosis efficiency is calculated as the exocytosis per unit Ca^2+^ charge transfer. We calculated an exocytosis efficiency of approximately 6 fF/pC in adult WT mouse RBCs after a 100 ms depolarizing pulse at 33° C. In adult vertebrate auditory hair cells, which also form ribbon synapses, similar exocytosis efficiencies of 5 to 6 fF/pC have been reported after 100 ms pulses (Chen & von Gersdorff, 2019; Moser et al., 2020). Rod bipolar cell ribbon synapses operate via nanodomain coupling between Ca^2+^ channels and docked vesicles (Jarsky et al., 2010), whereas immature calyx of Held synapses contain conventional active zones that operate via Ca^2+^ microdomains where Ca^2+^ influx triggers vesicle exocytosis from greater distances (Kushmerick, Renden, & von Gersdorff, 2006). Nevertheless, exocytosis efficiency can still be as high as 16 to 20 fF/pC at immature calyx of Held axon terminals after a short 1 ms pulse. Further studies will be required to verify the relationship between TPBG and nanodomain coupling at RBC ribbon synapses.

### TPBG and synaptic ribbon proteins

Neurotransmitter release from synaptic ribbons can be simplified into three major steps: synaptic docking, Ca^2+^ influx and its detection by Ca^2+^ sensors, and Ca^2+^-dependent fusion of synaptic vesicles with the presynaptic membrane (Dieck & Brandstätter, 2006; Morgans, 2000). These are all potential points at which TPBG may be supporting efficient vesicle release from RBCs. Vesicle accumulation and transport in RBC synaptic terminals is dependent on F-actin polymerization and is regulated by PKCα (Berglund, Midorikawa, & Tachibana, 2002; Doussau & Augustine, 2000; Job & Lagnado, 1998; Minami, Berglund, Sakaba, Kohmoto, & Tachibana, 1998; Tachibana, 1999). We originally identified TPBG as a PKCα-dependent phosphoprotein (Wakeham et al., 2019); thus, it is possible that PKCα controls vesicle docking via actin dynamics in RBC axon terminals through TPBG.

The tight temporal control of vesicle release that is a trademark of synaptic ribbons requires nanodomain-coupling between L-type voltage-gated Ca^2+^ channels and Ca^2+^ binding proteins that trigger synaptic vesicle fusion (Balakrishnan et al., 2015; Eggermann, Bucurenciu, Goswami, & Jonas, 2012; Jarsky et al., 2010). Synaptic ribbons are primarily composed of the scaffold protein RIBEYE and are anchored to the presynaptic active zone by a number of cytoskeletal anchoring proteins, including Bassoon and Piccolo (Zanazzi & Matthews, 2009). We did not observe a difference in RIBEYE immunofluorescence in WT and TPBG-KO axon terminals, but immunofluorescence does not have the resolution to confirm the correct localization or orientation of the ribbon at the active zone, If TPBG is required for correct localization of the ribbon, exocytosis efficiency may be significantly disrupted in KO RBCs, but electron microscopy showed normal positioning of ribbons at RBC active zones in the KO. Ca_v_1.4 and RIBEYE appear co-localized by immunofluorescence in both WT and TPBG-KO RBC axon terminals, but a small disruption of this tight coupling might not be detected with immunofluorescence and could have a large effect on glutamate release by slowing the accumulation of Ca^2+^ at the sensor. Nanodomain coupling likely depends on a close spatial relationship between the L-type channels and the vesicle release machinery, yet it is not fully known how this clustering is achieved in RBCs. It is possible that TPBG facilitates Ca^2+^ channel clustering or association between the channels and the ribbon such that, in its absence, normal nanodomain coupling is disrupted and synaptic vesicle fusion is reduced.

Ultrastructural analysis of RBC axon terminals revealed that the synaptic ribbons are approximately 15% smaller in TPBG-KO RBCs compared to WT. Shorter ribbons could result in a smaller number of ribbon-attached vesicles that may constitute a smaller ready releasable pool (RRP), resulting in the observed C_m_ jump attenuation. The density of ribbons was not reduced in the TPBG-KO as assayed by immunofluorescence imaging, nor were Ca^2+^ currents changed; thus, the smaller C_m_ jumps not could be due to a smaller number of ribbons or reduced Ca^2+^ influx. We used a large 100 ms depolarizing pulse to evoke vesicle release which likely exhausts the fast RRP (docked vesicles) and potentially depletes the secondary RRP (vesicles attached to the ribbon in upper rows). The fast RRP probably consists of the first row of vesicles lining the bottom of the ribbon closest to the active zone (von Gersdorff & Matthews, 1997; von Gersdorff et al., 1996) and this pool may not be affected by a shorter ribbon. In contrast, a shorter ribbon may significantly reduce the size of the secondary RRP, presumably corresponding to vesicles tethered higher on the ribbon, leading to reduced C_m_ jumps. Indeed, in RIBEYE-KO mice, where the synaptic ribbon is absent, exocytosis is greatly reduced in RBC and rod photoreceptor synapses (Grabner & Moser, 2021; Maxeiner, Luo, Tan, Schmitz, & Südhof, 2016). More experimentation is required to determine exactly how the shorter ribbons in the TPBG-KO RBC axon terminals affects the different vesicle pools.

### TPBG and reciprocal feedback inhibition

To confirm that the TPBG-KO reduces RBC vesicle fusion and glutamate release, we compared GABAergic feedback inhibition between WT and TPBG-KO RBCs. We found attenuation of inhibitory charge transfer in the TPBG-KO. With glycinergic inhibition blocked by strychnine, the primary remaining source of feedback inhibition in RBCs is A17-mediated GABAergic reciprocal feedback. GABA released from A17-ACs is sensed by two kinetically distinct ionotropic GABA receptors in RBC axon terminals: GABA_A_ and GABA_A_ρ (Fletcher et al., 1998). GABA_A_ receptor-mediated current is rapidly-activating, while GABA_A_ρ-mediated current is slowly-activating (Grimes et al., 2015). We detected two different inhibitory components: a large, quickly-inactivating current and a smaller sustained one. Though we did not pharmacologically isolate each current, the kinetics of the inhibitory currents suggest that peak I_Inhib_ is likely mediated by GABA_A_ receptors and sustained I_Inhib_ is likely mediated by GABA_A_ρ receptors. We note that our peak WT GABA_A_ receptor currents of ∼125 pA and sustained GABA_A_ρ receptor currents of ∼50 pA are considerably larger than reported previously in mouse RBCs (Chávez et al., 2010). Again, we attribute this to our use of older animals and higher recording temperatures and, perhaps, the use of Ames recording solutions: Ames contains ascorbic acid, which is an enhancer of GABA currents (Calero et al., 2011).

GABAergic reciprocal feedback from A17-ACs might be altered if GABA receptor expression or localization was disrupted by TPBG-KO. GABA_A_ receptors are large pentameric ion channels usually containing α, β, and γ subunits. Early work indicates that RBC axon terminals express GABA_A_α1, α3, and γ2 receptor subunits (Fletcher et al., 1998), but a recent study found that the α1 subunit dominates in adult mice (Sinha et al., 2021). GABA_A_ρ receptors contain only ρ subunits (Fletcher et al., 1998). Therefore, we used antibodies against GABA_A_α1 and GABA_A_ρ1 to visualize GABA_A_ and GABA_A_ρ receptor localization and an antibody against PKCα to isolate RBC-specific axon terminals. We did not detect any difference in GABA receptor immunofluorescence in RBC axon terminals between WT and TPBG-KO, indicating that reduced GABAergic feedback is likely not due to disrupted GABA receptor localization.

We found that TPBG-KO increased scotopic ERG b-waves, suggesting an increase in RBC depolarization. The scotopic ERG b-wave is initiated by current through TRPM1 non-specific cation channels in the RBC dendrites. Yet, we did not find any difference in CPPG-induced TRPM1 currents between WT and TPBG-KO retinas and the input resistance and excitability of the RBC was unchanged. However, the scotopic b-wave is also shaped by reciprocal inhibition. Pharmacological studies show that blocking inhibitory currents from third-order neurons changes the scotopic b-wave. In particular, injections of a GABA_A_ antagonist (bicuculline) or a GABA_A_ρ antagonist (TPMPA) into the IPL significantly increases the amplitude and duration of the scotopic b-wave (Dong & Hare, 2000, 2003), suggesting that GABAergic feedback from A17-ACs contributes to the shape of the RBC light response. We suggest that the enlarged ERG b-wave in the TPBG-KO is due to reduced inhibitory feedback onto the RBC terminals, secondary to reduced RBC glutamate release by the RBC.

## Summary

Our findings show that genetic deletion of TPBG attenuates scotopic ERG oscillatory potentials, synaptic vesicle exocytosis from RBCs, and reciprocal feedback onto RBCs. We propose that TPBG is required for efficient coupling of Ca^2+^ influx and glutamate release at RBC active zones, possibly through its effect on synaptic ribbons. Further evaluation of TPBG is critical for producing a more comprehensive understanding of its role in fine-tuning RBC synaptic ribbon physiology.

## Acknowledgements

This work was supported by NIH grants to CWM (R01 EY029985 and P30 EY010572) and HvG (R01 EY014043 and DC012938). The authors would like to thank R Lane Brown, Marc A Meadows, and André A Dagostin for assistance with patch clamp electrophysiology and data analysis. We would also like to thank the OHSU Neuroscience Imaging Center’s Ultrastructure and Single Particle Microscopy Core (P30 NS061800) for FIB-SEM data collection.

---

**Figure 2 Supplementary Figure:**
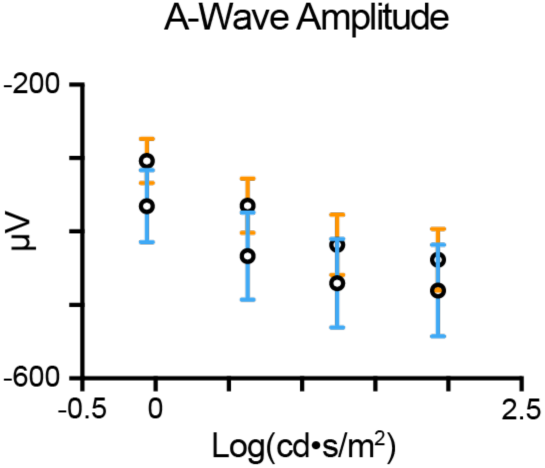
TPBG-KO does not alter the scotopic ERG a-wave. Quantification of a-wave amplitudes recorded after light stimuli (-0.06 to 1.93 log(cd•s/m^2^)). A-waves are measured from baseline to the a-wave trough. Data is presented as the mean ± SEM.

**Figure 4 Supplementary Figure:**
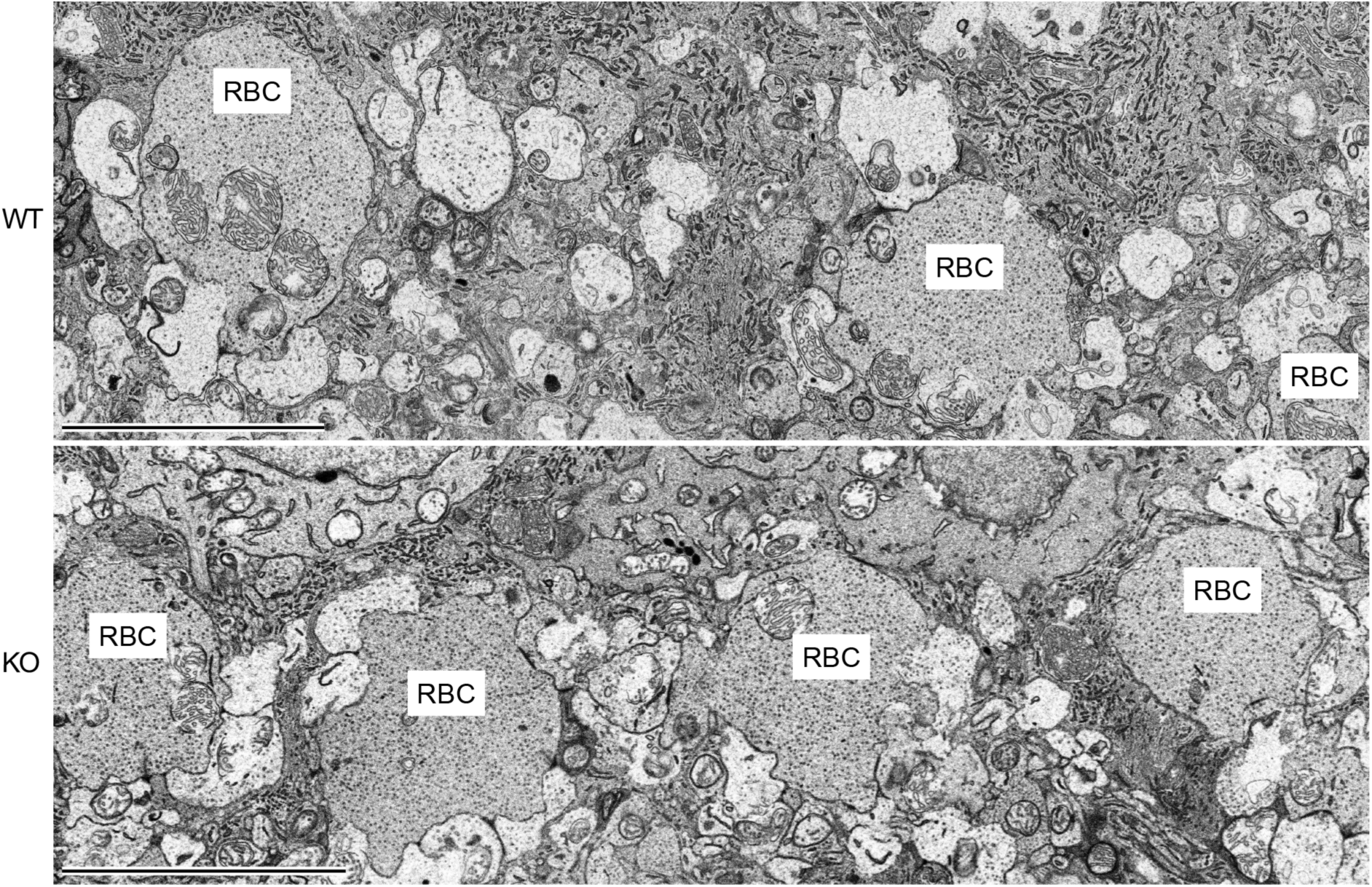
FIB-SEM imaging of RBC axon terminals. Focused ion beam scanning electron microscopy (FIB-SEM) was used to image RBC axon terminals in the distal IPL and examine synaptic ribbon location and morphology. Zoomed-out examples of WT and TPBG-KO (KO) RBC axon terminals in the IPL. The scale bar is 5 μm.

**Figure 5 Supplementary Figure:**
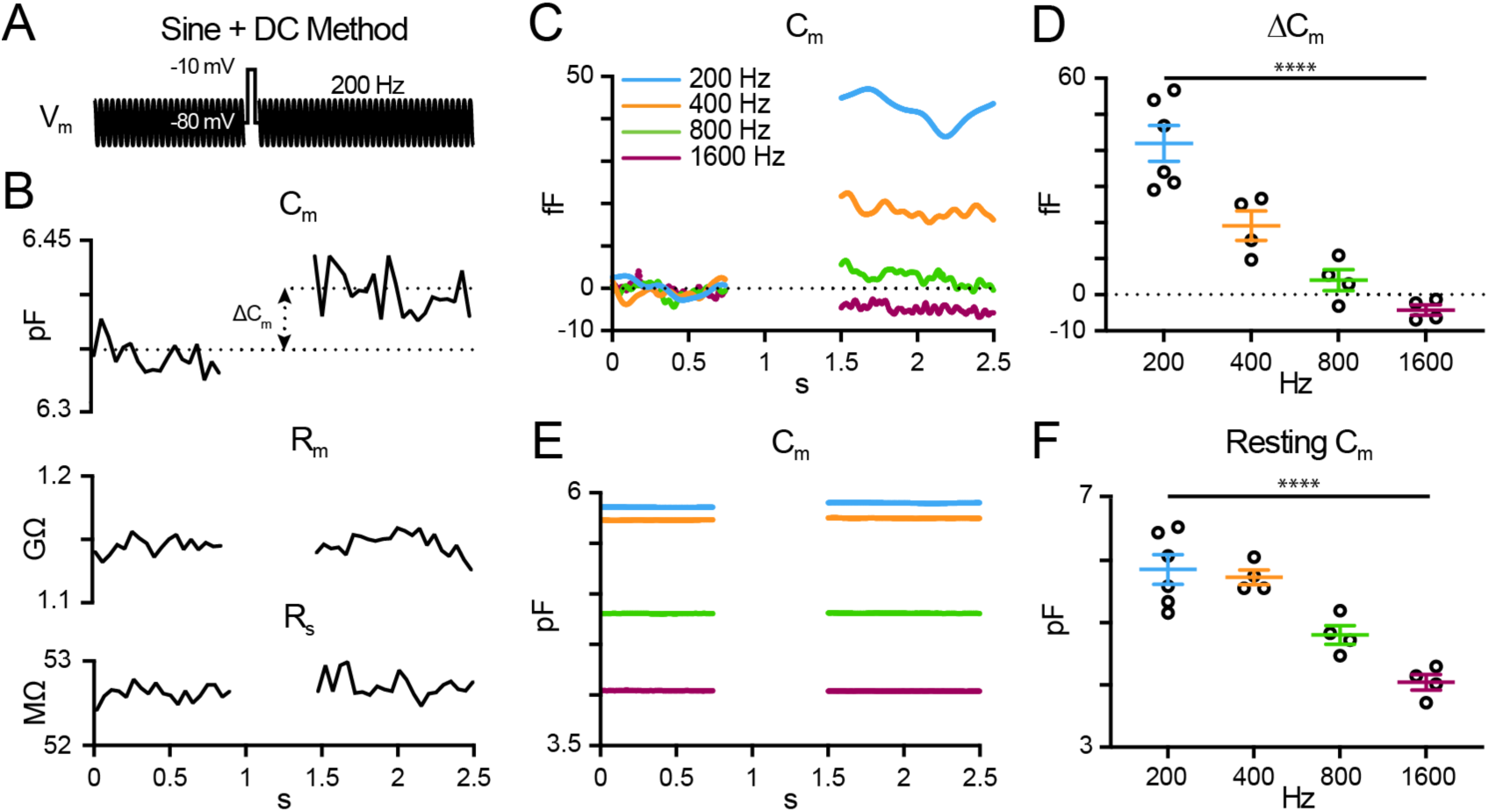
Measuring membrane capacitance in RBCs. **(A)** Whole-cell voltage-clamp capacitance recordings were made using the “sine + DC” method. A sinusoidal voltage command (200 Hz at 30 mV trough-to-peak) was superimposed on the holding potential of -80 mV. Membrane capacitance (C_m_), membrane resistance (R_m_), and series resistance (R_s_) were recorded before and after a depolarizing pulse from -80 mV to -10 mV for 100 ms. **(B)** A sample recording from a WT RBC using the sine + DC method as described. The change in membrane capacitance (ΔC_m_) was measured as the difference between the C_m_ before and after the depolarizing pulse. Changes to R_m_ (middle) and R_s_ (bottom) were small and do not correlate with changes in C_m_. **(C)** Average baseline-subtracted C_m_ recordings using 200 Hz (blue; *n* = 6), 400 Hz (orange; *n* = 4), 800 Hz (green; *n* = 4), or 1600 Hz (magenta; *n* = 4) sinusoidal voltage commands in WT RBCs. Data for the 200 Hz condition was reused from WT in Figure 4. Traces with higher frequencies were resampled to 200 Hz for ease of comparisons. **(D)** Quantification of ΔC_m_. **(E)** Average C_m_ recordings and **(F)** quantification of resting C_m_ across frequencies. Resting C_m_ was measured before the depolarizing pulse. Open circles in bar graphs represent single cells and bars represent the mean ± SEM. Statistical significance was determined using Brown-Forsythe ANOVA tests; ****: *p* < 0.0001.

